# Co-targeting KRAS and BCL-2 induces pyroptosis and antitumor tertiary lymphoid structures

**DOI:** 10.64898/2026.10.12.756217

**Authors:** Xu Yang, Man Li, Jian Zhang, Danming He, Xu Liu, Lihui Liu, Jianchun Duan, Zhijie Wang, Hua Bai, Jie Wang

**Affiliations:** State Key Laboratory of Molecular Oncology, CAMS Key Laboratory of Translational Research on Lung Cancer, Department of Medical Oncology, National Cancer Center/National Clinical Research Center for Cancer/Cancer Hospital, Chinese Academy of Medical Sciences and Peking Union Medical College, Beijing, 100021, China; National Cancer Center/National Clinical Research Center for Cancer/Cancer Hospital & Shenzhen Hospital, Chinese Academy of Medical Sciences and Peking Union Medical College, Shenzhen, 518116, China; Beijing Chest Hospital, Capital Medical University & Beijing Tuberculosisand Thoracic Tumor Research Institute

**Keywords:** KRAS G12D, mitochondrial vulnerability, pyroptosis, tertiary lymphoid structures

## Abstract

Mutant KRAS drives lung tumorigenesis through both cell-intrinsic and cell-extrinsic mechanisms, yet mutation subtype-specific differences in metabolic regulation and immune evasion remain incompletely understood. Here we show that the KRAS G12D mutation, particularly in the KRAS/STK11 co-mutant (KL) context, is associated with poor prognosis and a suppressed immune microenvironment characterized by impaired antigen presentation. At the cell-intrinsic level, G12D exhibits weakened interaction with hexokinase 1 (HK1), reduced spare respiratory capacity, and diminished mitochondrial stress resistance compared with G12C. Based on this mitochondrial vulnerability, we screened a panel of clinically available agents that induce mitochondrial stress and identified that co-targeting KRAS and BCL-2 synergistically triggers GSDME-mediated pyroptosis predominantly in G12D tumors. This combination further induces functional tertiary lymphoid structures (TLS) in vivo. Immune cell depletion experiments further demonstrate that the antitumor efficacy is dramatically dependent on B cells. Moreover, this treatment strategy induced membrane attack complex (MAC) deposition in situ and elicited memory antitumor responses. Collectively, these findings establish that combined KRAS and BCL-2 inhibition elicits pyroptosis and TLS formation, leading to B-cell-dependent in situ and memory antitumor immunity, providing a mechanistic rationale for targeting mitochondrial-immune crosstalk in KRAS G12D-mutant lung cancer.

## Introduction

KRAS is a major driver oncogene in lung cancer[1], with G12C, G12D, and G12V as the predominant mutation subtypes[2]. Co-occurring mutations or dysfunction in STK11 or TP53 further diversify the molecular and clinical landscape of KRAS-mutant tumors[3].

Historically, oncogenic KRAS mutations have been treated as functionally equivalent driver events. However, accumulating biochemical, structural, and genetic evidence has fundamentally challenged this paradigm. Different amino acid substitutions at the same codon—and across different codons—confer distinct molecular properties. For instance, Q61L/H mutations severely impair intrinsic GTP hydrolysis (by 40- to 80-fold), while G13D uniquely accelerates nucleotide exchange rates by approximately 10- to 14-fold relative to wild-type[4, 5]. These biochemical distinctions translate into differential effector engagement. G12D, for instance, is predicted to drive the lowest RAF activation among common KRAS mutants due to its weak RAF binding and rapid intrinsic hydrolysis[4]. Consequently, individual KRAS alleles may display preferential engagement of downstream pathways.

Emerging evidence further indicates that these signaling biases drive allele-specific metabolic reprogramming, tumor phenotypes, and immune microenvironments. Metabolomic profiling of isogenic colorectal cancer cell lines reveals that G12D, G12V, and G13D mutants each exhibit distinct metabolic signatures involving glycolysis, glutamine utilization, and nucleotide metabolism[6]. Moreover, the KRAS4A splice variant directly interacts with hexokinase 1 on the mitochondrial outer membrane to enhance glycolytic flux, a function not shared by KRAS4B[7, 8]. The complexity of KRAS-driven oncogenesis is further amplified by co-occurring genetic alterations, which shape the biological consequences of specific KRAS alleles in a tissue-dependent manner. Large-scale genomic analyses have uncovered allele-specific co-mutation patterns that define major subsets of KRAS-mutant tumors[3, 9]. Importantly, these co-mutations are not passive bystanders but actively modulate tumor behavior and therapeutic response. STK11 loss, for instance, promotes neutrophil recruitment and T-cell exclusion, conferring primary resistance to PD-1/PD-L1 blockade specifically in the context of KRAS-mutant lung adenocarcinoma[10, 11]. These observations underscore that the functional impact of a given KRAS mutation cannot be interpreted in isolation—it is profoundly modified by the concurrent mutational landscape and tissue context.

Here, we identified a selective mitochondrial vulnerability in G12D through systematic analysis integrating mutation subtypes and co-mutation status. Exploiting this weakness via a focused screen, we found that co-targeting KRAS and BCL-2 synergistically triggers GSDME-mediated pyroptosis predominantly in G12D tumors, which further induces tertiary lymphoid structures and B cell dependent antitumor immunity. These findings establish a strategy to convert allele-specific biological differences into targeted therapeutic opportunities for KRAS G12D mutant lung cancer.

## RESULTS

### G12D mutation is associated with poor prognosis and immune suppression in KL tumors

First, we assessed overall survival across distinct KRAS mutation and co-mutation subtypes in a published pan-cancer cohort[12]. In KRAS-mutant lung cancers, patients harboring G12D mutation exhibited worse overall survival than those with G12C or non-G12D mutations (Figure 1a and sFigure 1a). This association was observed exclusively in KL (KRAS-STK11 comutation) lung cancers and was absent in KP (KRAS-TP53 comutation)and KC (KRAS-CDKN2A comutation) subtypes (sFigure 1a). We recapitulated these findings in an independent published cohort of patients with lung cancer who received immunotherapy[13] (Figure 1a and sFigure 1b). To further validate our results, we compared progression-free survival (PFS) and treatment response in our single-center cohort. Among KL patients, those with the G12D mutation achieved worse PFS and diminished treatment response (Figure 1a). We subsequently compared the microenviroment among mutation and co-mutation subtypes using TCGA and GSE72094[14] datasets. G12D mutation was associated with lower ESTIMAT and IMMUE scores in patients with STK11 co-mutation or low expression given that KL tumor featuring low STK11 expression[3] (Figure 1b and sFigure 1c). Further IPS analysis revealed that G12D mutation was associated with antigen presentation deficits (Figure 1b), marked by reduced effector cells and elevated suppressor cells (sFigure 1c). To further dissect the cellular composition of the tumor microenvironment in a genetically defined context, we crossed Kras^G12D/+ and Kras^G12C/+ mice with Stk11^f/f transgenic mice to generate Kras^G12D/+;Stk11^f/f and Kras^G12C/+;Stk11^f/f genotypes. Kras^G12D/+;Stk11^f/f mice exhibited significantly increased lung weight, indicating enhanced tumorigenicity (Figure 1c). Single-cell analysis revealed that Kras^G12D/+;Stk11^f/f tumors were characterized by reduced CD8+ T cells and increased macrophages and dendritic cells (DCs) (Figure 1d and sFigure 1d). GSEA analysis further demonstrated that macrophages and DCs exhibited attenuated antigen presentation and interferon response functions (Figure 1e and 1f), consistent with human tissue data. Finally, immunohistochemistry (IHC) analysis confirmed that CD8+ T cell infiltration was decreased in KL lung cancer patients with STK11 mutation or low expression compared with other subtypes (Figure 1g). Together, these findings suggested that the unfavorable prognosis of G12D mutant patients was associated with an immunosuppressive microenvironment characterized by myeloid cell dysfunction.

**Figure 1.**
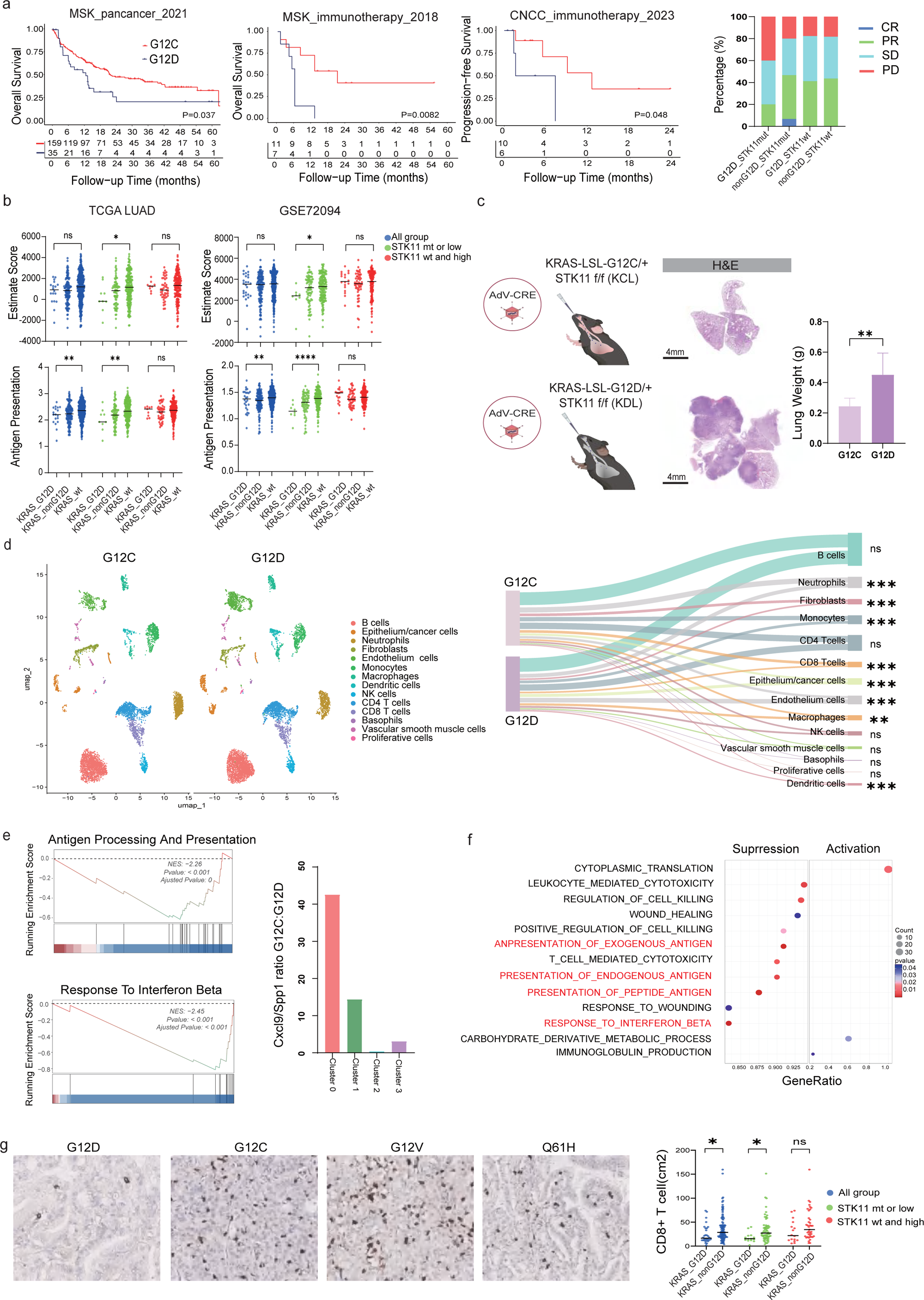
KRAS G12D is associated with poor prognosis and an immunosuppressive tumor microenvironment in KL lung cancers. a, Kaplan–Meier survival curves for overall survival (OS) in a published pan-cancer cohort (left) and an immunotherapy cohort (middle), stratified by KRAS mutation and co-mutation subtypes; progression-free survival (PFS) and treatment response in the single-center cohort (right). b, IPS (up) and ESTIMATE score analysis (down) from TCGA (left) and GSE72094 (right) datasets comparing G12D versus non-G12D tumors within STK11-low/co-mutant KL settings. c, Lung weights of Kras^G12D/+;Stk11^f/f (KDL) versus Kras^G12C/+;Stk11^f/f (KCL) mice. d, Single-cell RNA-seq showing proportions of CD8+ T cells, macrophages, and dendritic cells (DCs) in KDL versus KCL tumors. e, f, GSEA of antigen presentation and interferon response pathways in macrophages (e) and DCs (f) from single-cell data. g, Representative IHC images of CD8+ T cell infiltration (left) and quantitative analysis (right) in KL patients with STK11 mutation/low expression versus other subtypes. Data are shown as mean ± SEM. *P < 0.05; **P < 0.01; ***P < 0.001.

### KRAS G12D subtype silences the interferon axis through reduced mtDNA leakage

To further explore the cell-intrinsic properties of KRAS G12D, we integrated TCGA, GSE72094, and CCLE datasets for KL lung cancers and observed a co-downregulation of interferon α response, interferon γ response, and inflammatory response pathways in G12D-mutant tumors (Figure 2a and sFigure 1e). Mechanistically, IFN signaling has been broadly reported to be associated with cytosolic DNA sensing pathways, particularly those involving mitochondria-derived DNA (mtDNA) [15]. This pathway is further interconnected with apoptosis[16], p53 signaling[17], and nuclear factor kappaB (NF-kappaB)[18] through mitochondrial and genomic stress (Figure 2b). GSVA analysis performed on KL lung cancer cell lines from the CCLE dataset confirmed the anticipated signature spectrum (Figure 2b). Subsequently, GSEA of single-cell RNA sequencing data from transgenic mice validated these findings, demonstrating downregulation of the apoptosis, p53, hypoxia, and TNFα via NF-kappaB signaling pathways in Kras^G12D/+;Stk11^f/f tumors (Figure 2c, 2d and sFigure 1f). Next, qPCR and cell culture supernatant ELISA experiments validated that IFN and the classic interferon-stimulated gene (ISG) CXCL10 were downregulated in the A427-G12D cell line (Figure 2e). We further assessed mitochondrial membrane potential (MMP) using JC-1 dye detected by confocal microscopy and flow cytometry. Indeed, G12D cell lines exhibited higher MMP than other mutations, regardless of STK11 mutation status (Figure 2f and 2g). To further exclude the possible influence of STK11 function on these results[19], we knocked out STK11 in KP cell lines and over-expressed STK11 in KL cell lines, respectively (sFigure 2a). G12D cells consistently exhibited elevated MMP under these conditions, further excluding a potential confounding role of STK11 (sFigure 2c-2f). Next, we generated KDL and KCL organoids from Kras^G12D/+;Stk11^f/f and Kras^G12C/+;Stk11^f/f mice, respectively. KDL organoids again showed higher MMP than KCL organoids (Figure 2h). To further investigate mitochondrial morphology, we employed super-resolution live-cell imaging (OMX SR). G12D mutant cells exhibited significantly elongated mitochondria, whereas G12C mutant cells displayed a punctate morphology, further supporting a distinct mitochondrial homeostasis phenotype associated with G12D (Figure 2i). As cytosolic mtDNA functions as a key link between mitochondrial status and IFN signaling, we detected cytosolic mtDNA by confocal microscopy and qPCR. As expected, less mtDNA was detected in G12D mutant cell lines (Figure 2j). Notably, nuclear-derived DNA accounted for only a minimal fraction of the cytosolic DNA pool, indicating that mtDNA was the primary contributor to IFN signaling in this context (Figure 2j). Finally, depletion of mtDNA using ethidium bromide (EtBr) or treatment with a STING inhibitor markedly decreased CXCL10 levels in the supernatant of G12C cells, abolishing the difference between G12C and G12D cells (Figure 2k). Collectively, these findings establish that STING-dependent sensing of mtDNA underlies the distinct IFN signaling phenotypes between G12D and G12C mutations.

**Figure 2.**
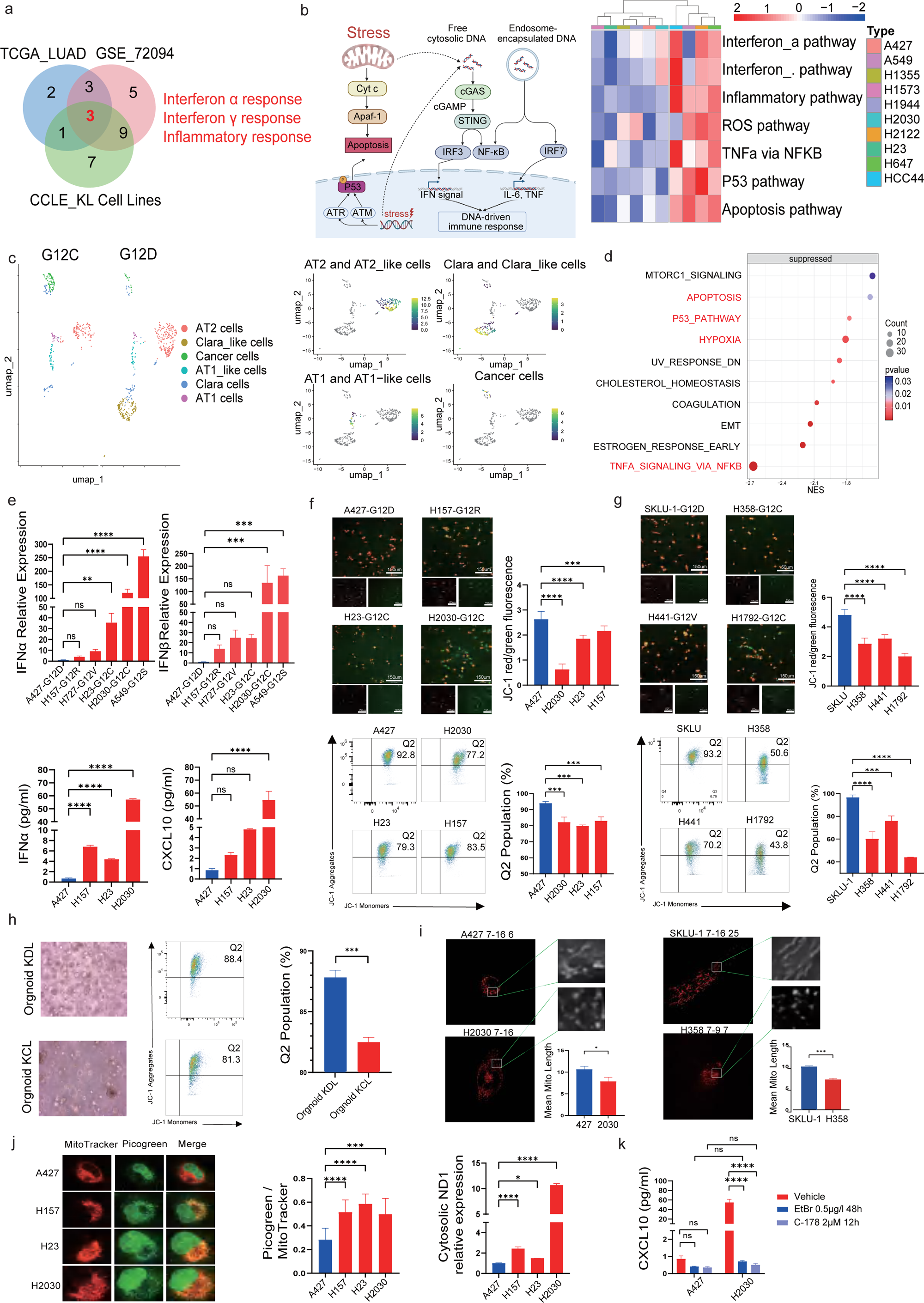
KRAS G12D silences the interferon axis by limiting mtDNA leakage. a, Venn diagram shew downregulated pathways in KRAS G12D versus non-G12D KL tumors from the TCGA (n = 86), GSE72094 (n = 78), and CCLE (n = 10) datasets based on GSEA. b, Schematic diagram illustrated the proposed crosstalk between IFN signaling, cytosolic mtDNA sensing, and stress-associated pathways. c, d, GSEA of single-cell RNA sequencing data from Kras^G12D/+;Stk11^f/f and Kras^G12C/+;Stk11^f/f tumors. e, qPCR (up) and ELISA (down) analysis of IFN and CXCL10 expression in human KL cell lines. n = 3 biological replicates, each with 3 technical replicates. f, g, Representative confocal microscopy images (up) of JC-1 staining. Red fluorescence indicates JC-1 aggregates; green fluorescence indicates JC-1 monomers; Quantification of MMP by flow cytometry (down) in human cell lines. n = 3 biological replicates, each with 3 technical replicates. h, Quantification of MMP by flow cytometry in transgenic mouse-derived organoids. n = 3 biological replicates, each with 3 technical replicates. i, Super-resolution live-cell imaging of mitochondrial morphology in human cell lines. n = 3 biological replicates, with 10-27 cells analyzed per replicate. j, Representative confocal microscopy images of cytosolic DNA staining; Detection of cytosolic DNA by qPCR. n = 3 biological replicates, each with 3-4 technical replicates. k, ELISA quantification of CXCL10 in G12C and G12D cells after EtBr or STING inhibitor treatment. Statistical significance was determined by one-way ANOVA test. *P < 0.05; **P < 0.01; ***P < 0.001; ****P < 0.0001.

### KRAS G12D exhibits weakened HK1 interaction, reduced spare respiratory capacity, and impaired stress resistance

Having observed a relatively stable mitochondrial homeostasis phenotype in KRAS G12D cells, we next performed Seahorse analysis. G12D cells exhibited reduced spare respiratory capacity (SRC), whereas no consistent differences were observed in classic KRAS downstream signaling cascades, including MAPK, PI3K, and mTOR both in either KL or KP cell lines (Figure 3a), indicating that alternative, non-canonical pathways are involved in modulating mitochondrial respiration. Interaction between mutant KRAS and hexokinase, particularly HK1, via the RBD-like domain of HK1 modulates the hexokinase activity on the outer mitochondrial membrane[8]. Given that different KRAS mutation subtypes have been reported to harbor distinct affinities for the RBD[20], we hypothesized that differential binding affinity between KRAS mutation subtypes and HK1 contributes to the observed differences in mitochondrial respiration. Then, we transduced BEAS-2B cells with lentivirus expressing KRAS G12C or G12D. To elucidate the biological impact of KRAS protein localization across different mutation subtypes, we also overexpressed KRAS G12C186S and G12D186S to disrupt KRAS membrane anchoring. Consistent with our previous findings, G12D BEAS-2B cells exhibited reduced SRC and higher MMP than their G12C counterparts, whereas these differences were abrogated between G12D186S and G12C186S cells (Figure 3b-c and sFigure 3a). Co-IP experiments further revealed that G12D exhibited weaker interaction with HK1 than G12C (Figure 3d). Consistent with the mitochondrial phenotype, this difference in binding strength was eliminated upon disruption of KRAS membrane anchoring, as no significant difference was observed between G12D186S and G12C186S cells (Figure 3d). Next, we overexpressed HK1 in the aforementioned BEAS-2B cells. As expected, HK1 overexpression elevated SRC (Figure 3e), and decreased MMP (Figure 3f and sFigure 3b). Furthermore, HK1 co-IP again revealed that G12D exhibited reduced binding affinity for HK1 compared with G12C, whereas neither G12D186S nor G12C186S showed detectable binding to HK1 (Figure 3g).

**Figure 3.**
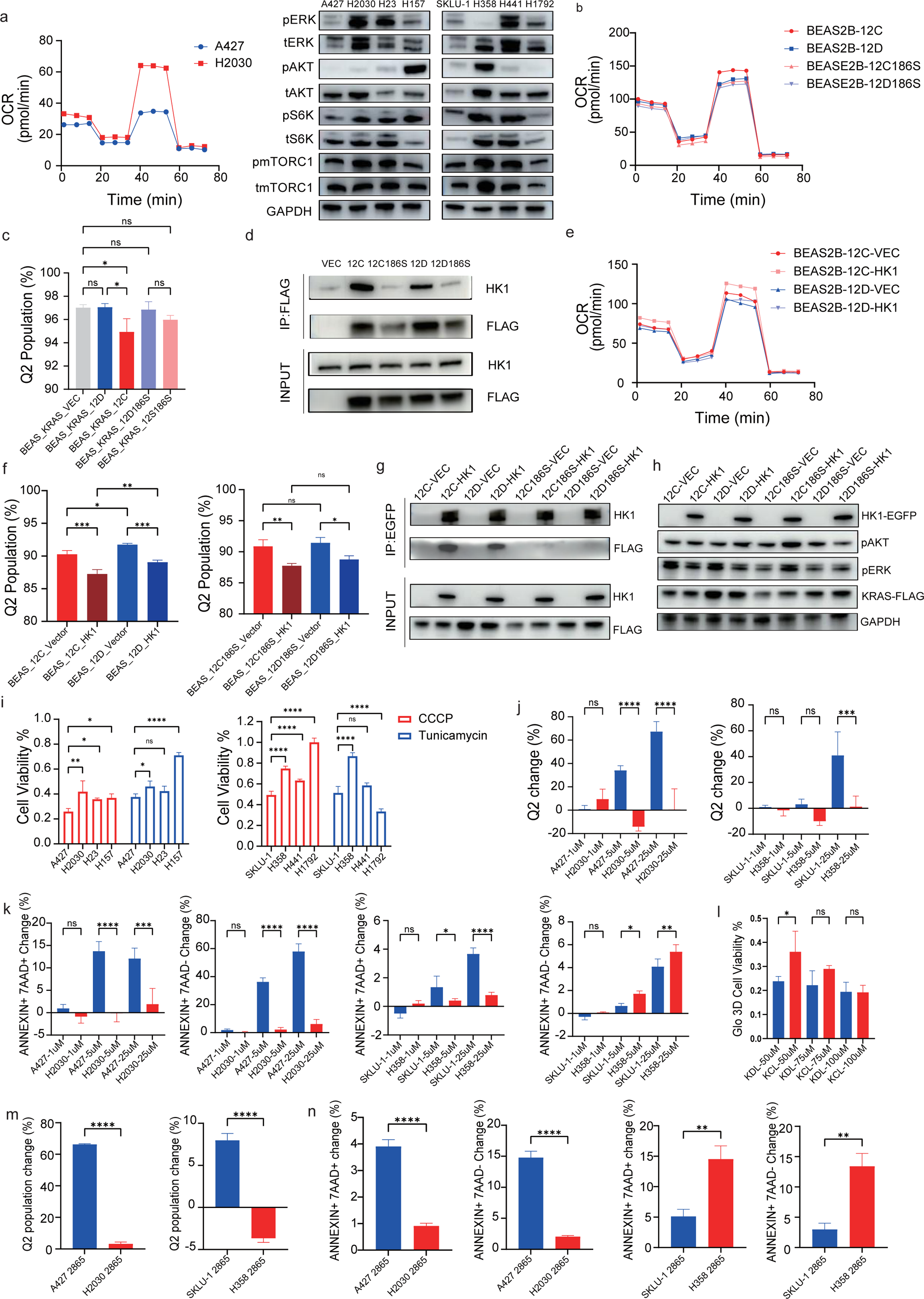
KRAS G12D exhibits altered HK1 interaction, mitochondrial respiration, and stress response. a, Seahorse measurement in KL cells and KRAS pathway immunoblots in KL and KP cells. b, c, Seahorse and MMP in BEAS-2B cells expressing FLAG-tagged KRAS G12C, G12D, G12C186S and G12D186S mutants. d, Co-immunoprecipitation (co-IP) analysis of KRAS-HK1 interaction in BEAS-2B cells expressing different KRAS mutants. e, f, Seahorse and MMP analysis in BEAS-2B cells expressing FLAG-tagged KRAS mutants with or without HK1-EGFP overexpression. g, co-IP analysis of HK1-KRAS interaction in BEAS-2B cells expressing different KRAS mutants with or without HK1-EGFP overexpression. h, KRAS pathway immunoblots in BEAS-2B cells expressing FLAG-tagged KRAS mutants with or without HK1-EGFP overexpression. i, Cell viability assessment by CCK8 assay in G12D and G12C cells following CCCP or tunicamycin treatment. j, k, MMP and apoptosis assays in G12D and G12C cells following CCCP treatment. l, CellTiter-Glo 3D viability assay in KDL and KCL organoids following CCCP treatment. m,n MMP and cell viability in G12D and G12C cells following BI-2865 treatment at IC50 concentration. n = 3 biological replicates, each with 3-4 technical replicates. Western blots and co-IP were representative of three independent biological replicates. Statistical significance was determined by one-way ANOVA test. *P < 0.05; **P < 0.01; ***P < 0.001; ****P < 0.0001.

Interestingly, HK1 overexpression led to decreased pERK levels in both G12C and G12D cells, but not in their membrane-anchoring-deficient counterparts (G12C186S and G12D186S) (Figure 3h). This observation further supports a competitive interaction between HK1 and the RAF-MEK-ERK pathway for KRAS binding, which is dependent on KRAS membrane localization. We next assessed mitochondrial stress sensitivity in G12D versus G12C cells, given the established link between enhanced spare respiratory capacity (SRC) and resistance to cell death[21]. Mechanistically, HK1 binding to the voltage-dependent anion channel (VDAC) promotes channel closure and prevents mitochondria-mediated apoptosis[22], while KRAS4A enhances HK1 oligomerization on the mitochondrial outer membrane[7]. As expected, G12D cells were vulnerable to CCCP but not to tunicamycin, an endoplasmic reticulum (ER) stress-inducing agent (Figure 3i). Further JC-1 and apoptosis assays revealed that G12D cells exhibited markedly decreased MMP (Figure 3j and sFigure 3c) and increased proportion of Annexin V+/7-AAD+ cells (Figure 3k and sFigure 3d). KDL organoids confirmed the selective vulnerability to mitochondrial stress observed in cell lines (Figure 3l). Given the interplay between KRAS signaling cascades and the anti-mitochondrial apoptosis pathway[23], as well as the observation that blocking downstream KRAS signaling induces mitochondrial apoptosis[24], we compared the treatment response of G12D and G12C cells to the pan-KRAS inhibitor BI-2865 at its IC50 concentration. Interestingly, G12D cells exhibited a more pronounced decrease in MMP (Figure 3m), yet this did not translate into increased cell death in both KL and KP cell lines (Figure 3n). To further dissect the cell signaling landscape in BI-2865-treated A427 cells, we isolated cells that did not exhibit a decrease in MMP following BI-2865 treatment by flow cytometry-based sorting. Bulk RNA analysis revealed that apoptosis pathway signature and pro-apoptotic proteins were not elevated in these treated cells with no MMP decrease (sFigure 3e), indicating that combination treatment targeting mitochondrial stress vulnerability is required to effectively eliminate this resistant subpopulation.

### Combined KRAS and BCL-2 inhibition induces pyroptosis in KRAS G12D tumors

BH3 mimetics[25], chemotherapy[26], and mitochondrial complex inhibitors[27] are known to induce mitochondrial stress and apoptosis in cancer. We therefore evaluated their synergetic effects with BI-2865 in A427 cells. Interestingly, the BCL-2 inhibitor, ABT-199, exhibited the highest synergetic score (Figure 4a and sFigure 4a). This synergistic effect was specific to the G12D mutation, as validated in transgenic mouse-derived organoids and human cell lines (Figure 4b and sFigure 4b). Flow cytometry experiments further revealed that the combination treatment dramatically reduced MMP (Figure 4c) and increased the proportion of Annexin V+/7-AAD+ cells in G12D-mutant human cells compared with their G12C counterparts (Figure 4d, 4e, sFigure 4c and sFigure 4d). This observation was recapitulated in murine KP-1 lung cancer and FC1242 pancreatic cancer cells (Figure 4f, 4g, sFigure 4e and sFigure 4f). As Annexin V+/7-AAD+ cells represent late-stage apoptotic or necrotic cells, we further examined cell morphology using phase-contrast microscopy. Remarkably, the combination treatment induced cell bubbling in G12D cells, which was absent in single-agent treatments (Figure 4d-4g). As this bubbling cell morphology has been broadly reported to be associated with pyroptosis[28, 29], we further examined the known GSDM family proteins. GSDME was markedly cleaved with elevated cleaved-caspase 3, whereas GSDMD-mediated pyroptosis, pMLKL-mediated necroptosis, and ACSL4-associated ferroptosis were not detected as the primary cell death modalities in A427 and KP-1 cell lines, as confirmed by immunoblotting and pharmacological inhibition assays (Figure 4h and sFigure 4g). We next examined the release of damage-associated molecular patterns (DAMPs) following combination treatment. Notably, the combination regimen significantly increased cell surface calreticulin exposure and HMGB1 release in the supernatant, indicative of immunogenic signals associated with pyroptosis (Figure 4i and sFigure 4h).

**Figure 4.**
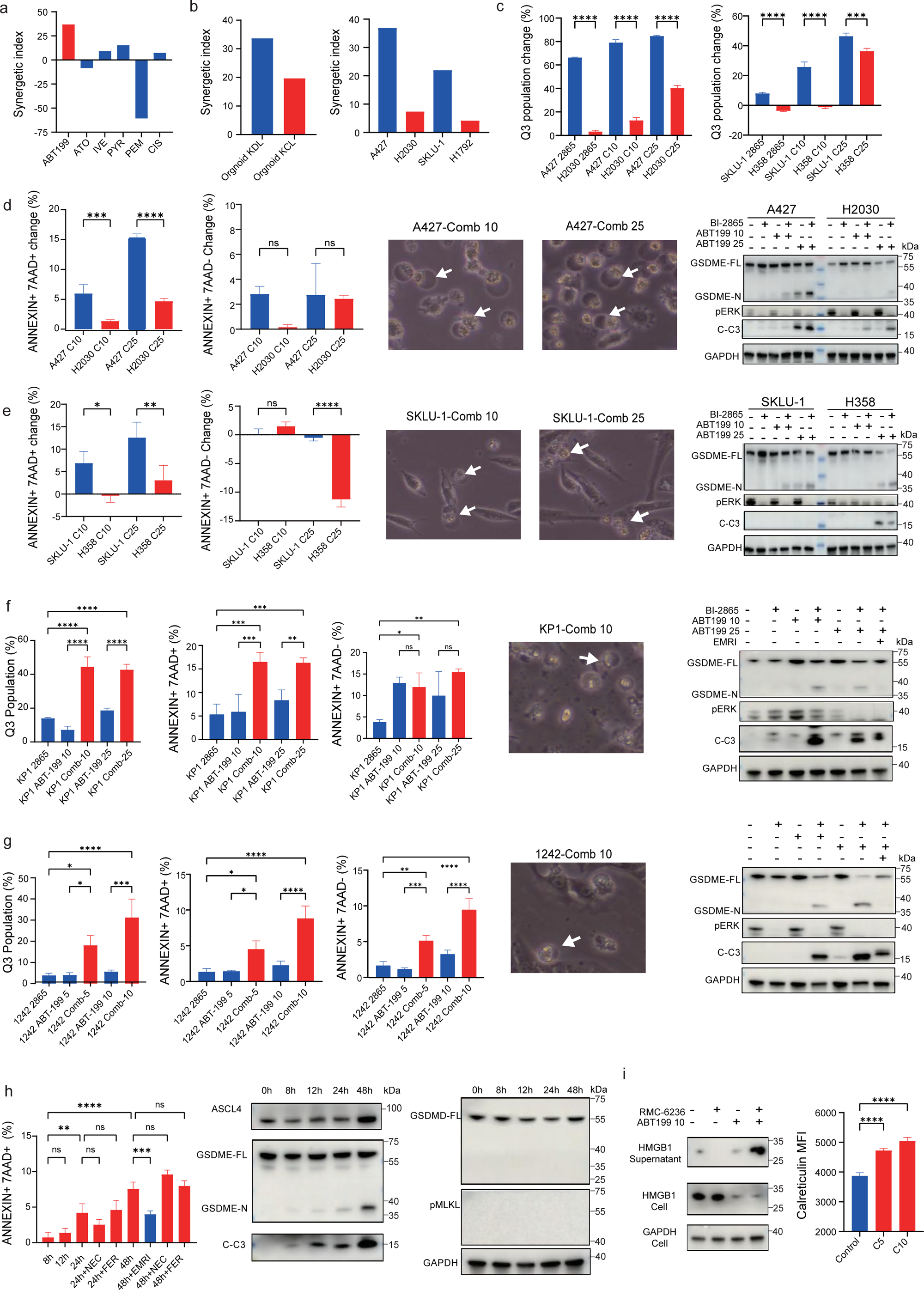
Pyroptosis induction by combined KRAS and BCL-2 inhibition in G12D cells in vitro. a, Synergistic effects of BI-2865 combined with BH3 mimetics, chemotherapeutic agents, or mitochondrial complex inhibitors in A427 cells. b, Synergistic effect of combined BI-2865 and ABT-199 in transgenic mouse-derived organoids and human cell lines. c, Flow cytometry analysis of MMP change following combination treatment. n = 3 biological replicates, each with 3 technical replicates. d, e, Flow cytometry, Phase-contrast microscopy images and immunoblot in human KL and KP cell lines. n = 3 biological replicates, each with 3 technical replicates. f, g, Flow cytometry, Phase-contrast microscopy images and immunoblot in mouse KP1 lung cancer and FC1242 pancreatic cancer cell lines. n = 3 biological replicates, each with 3 technical replicates. h, Flow cytometry of apoptosis and immunoblot of cell death markers in A427 cell lines following combination treatment. n = 3 biological replicates, each with 3 technical replicates. I, Cell supernatant HMGB1 (left) and Surface calreticulin (right) in A427 cell lines following treatment. n = 3 biological replicates, each with 3 technical replicates. Western blots were representative of three independent biological replicates. Statistical significance was determined by one-way ANOVA test. *P < 0.05; **P < 0.01; ***P < 0.001; ****P < 0.0001.

### KRAS and BCL-2 inhibitors confer a synergistic effect in vivo

We next evaluated the antitumor efficacy of the combination treatment in vivo. This regimen demonstrated superior antitumor efficacy and prolonged overall survival in both KP-1 and FC1242 subcutaneous xenograft models (Figure 5a, 5b, sFigure 5a and sFigure 5b). Flow cytometry analysis revealed a marked expansion of CD45+ immune cells in the combination group. However, neither the proportions of CD4+ or CD8+ T cells among CD45+ cells, nor the frequencies of CD44+CD4+ or CD44+CD8+ T cells, were significantly altered. Furthermore, no increase was observed in the percentages of GzmB+ or IFNy+ CD8+ T cells among live cells. Strikingly, the combination treatment led to a substantial increase in both CD3-cells and CD19+ B cells, accompanied by a corresponding decrease in the relative abundance of CD3+ T cells. The reduction in CD3+ T cells was consistent with a previous study reporting that BRAF and MEK inhibitors regulate the tumor immune microenvironment via pyroptosis[30]. Alongside these changes, we observed an expansion of both total CD49b+ NK cells and their GzmB+ cytotoxic subset (Figure 5c and sFigure 5c). Among the myeloid compartment, the frequency of dendritic cells (DCs) and tumor-associated macrophages (TAMs) showed a modest decrease, while the proportions of myeloid-derived suppressor cells (MDSCs) remained comparable between groups. Functionally, however, we observed a significant increase in CD86+ and CD103+ DC subsets, indicative of enhanced DC maturation and antigen presentation. By contrast, the proportion of M1-polarized TAMs was not altered (Figure 5d and sFigure 5d). In summary, the combination treatment reshaped the tumor microenvironment characterized by abundant B cells and enhanced NK and DC functions.

**Figure 5.**
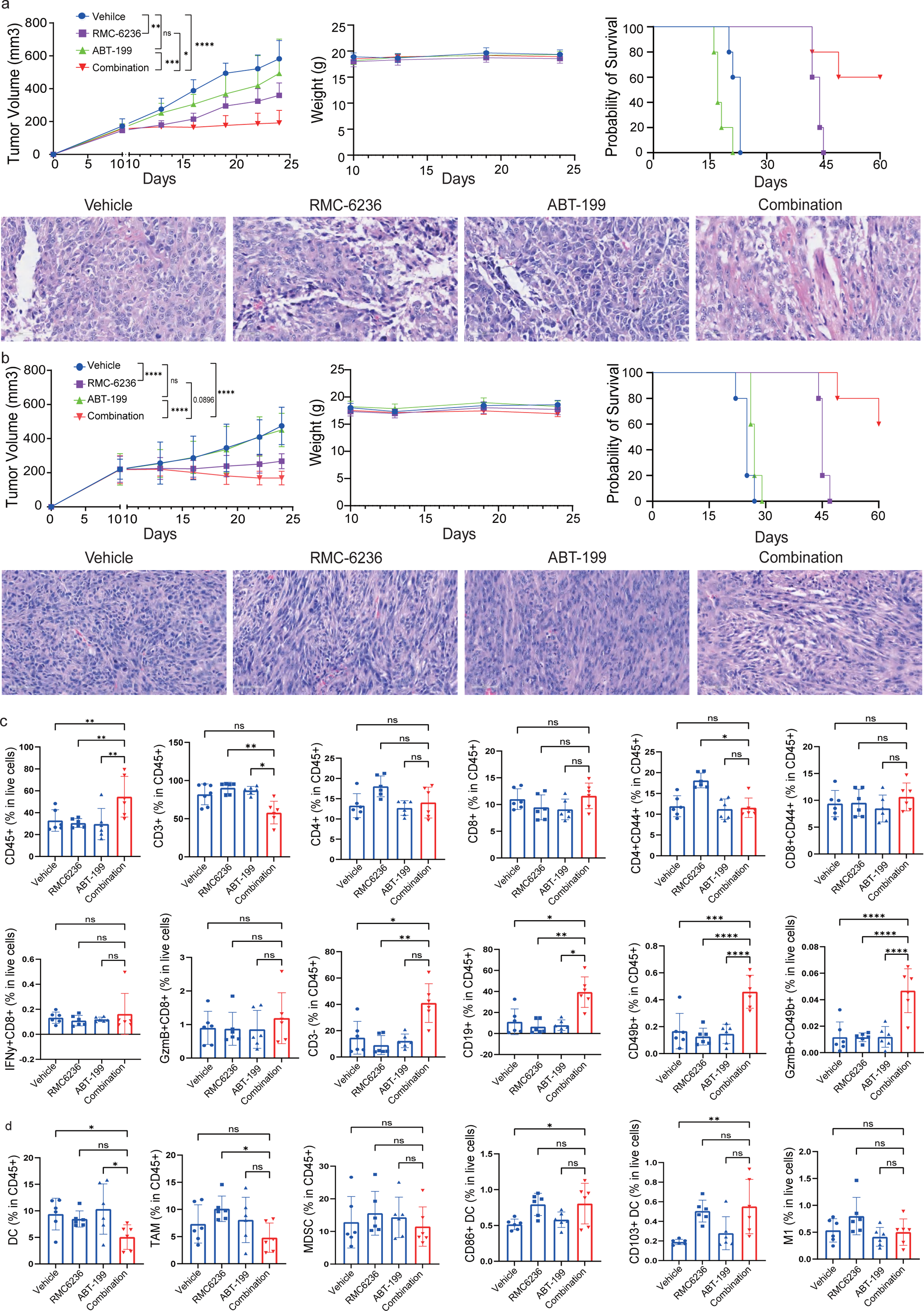
In vivo synergistic antitumor effect of combined KRAS and BCL-2 inhibition. a, b, Antitumor efficacy of combination treatment in KP-1 and FC1242 subcutaneous xenograft models. n = 8 mice per group. c, Lymphoid immune cell composition in KP1 tumors. n = 6 per group. d, Myeloid immune cell composition in KP1 tumors. n = 6 per group. Statistical significance was determined by one-way ANOVA test. *P < 0.05; **P < 0.01; ***P < 0.001; ****P < 0.0001.

### Combined KRAS and BCL-2 Inhibition Induces Tertiary Lymphoid Structures

To determine whether these B cells formed organized lymphoid aggregates, we next examined tumor sections by H&E staining. Notably, multiple B cell aggregates appeared in the combination group, whereas none were observed in vehicle or single-agent groups (Figure 6a). Multiplex immunofluorescence further revealed organized structures positive for CD3, CD19, CD23, and CD31, confirming the presence of tertiary lymphoid structures (TLS) (Figure 6b). In germinal centers, B cells undergo affinity maturation and class switching to produce high-affinity IgG antibodies, which function as key effector molecules in B-cell-mediated antitumor immunity[31, 32]. As expected, multiplex immunofluorescence revealed increased CD138+ plasma cells and an elevated IgG/IgA ratio following combination treatment (Figure 6c and 6d). The number of B cells increased markedly at D2 and D4, and declined thereafter (Figure 6e). BCR repertoire analysis revealed dynamic changes in clonal diversity. The Shannon and inverse Simpson indices were significantly elevated at D2 and D4, indicating progressive diversification of the B cell repertoire with concomitant contraction of dominant clones (Figure 6f). By D12, clonal diversity declined while dominant clones emerged, accompanied by an initial expansion followed by a reduction in clone count and somatic hypermutation (SHM) high clones (Figure 6f). This temporal trajectory—from clonal diversification to clonal selection—is highly consistent with the germinal center reaction. Furthermore, isotype analysis showed that IgG reached its peak at D12, coinciding with the maturation of the humoral response (Figure 6g). Together, these single-cell data provided molecular evidence for germinal center dynamics and class switching, supporting the maturation of B- cell-dependent antitumor immunity following combination treatment. Surprisingly, we also observed increased Shannon and inverse Simpson indices, along with elevated SHM, in the peripheral blood of treated mice (Figure 6h). These peripheral BCR features correlated positively with intratumoral TLS, suggesting a systemic imprint of local TLS formation.

**Figure 6.**
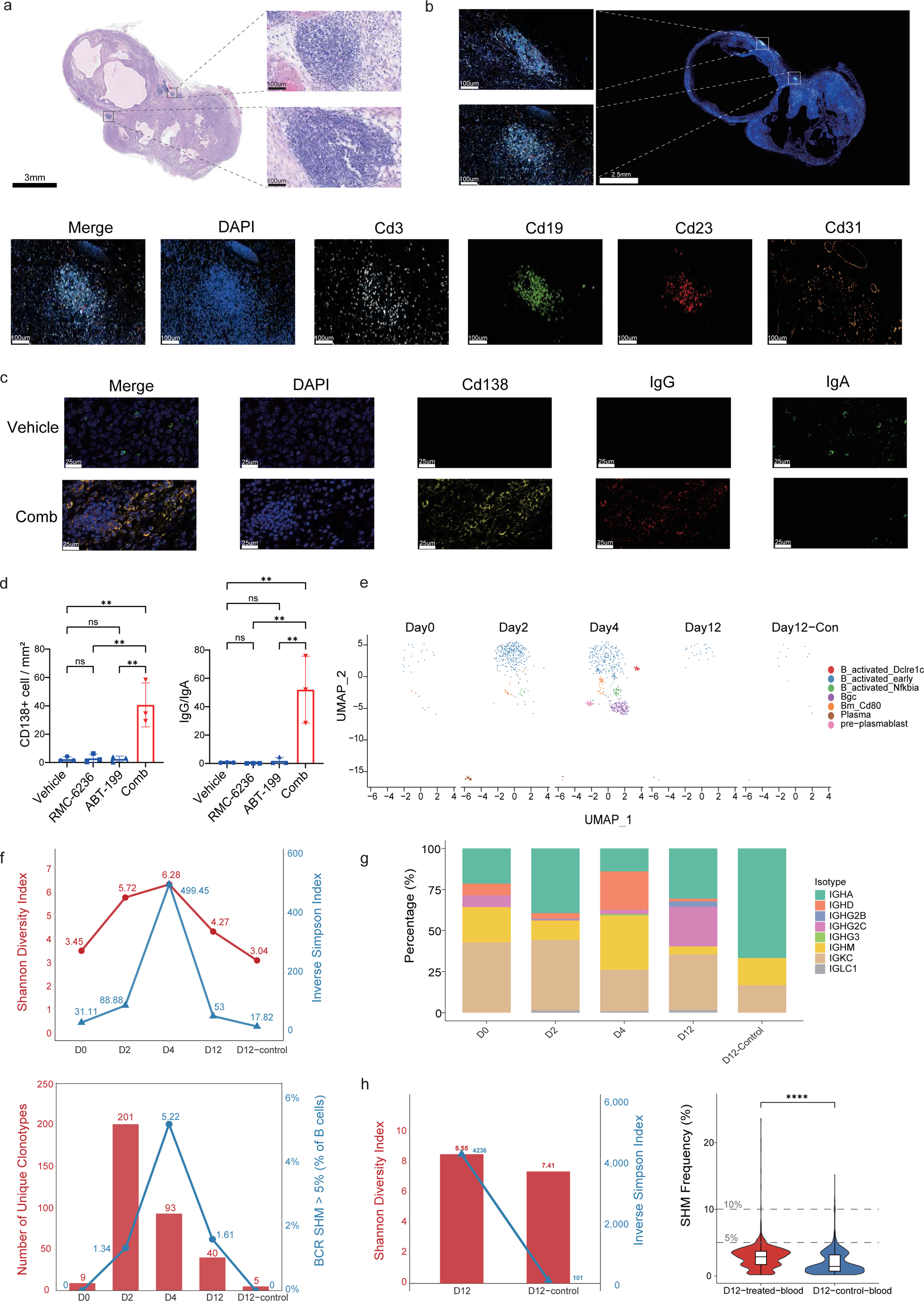
Combined KRAS and BCL-2 inhibition induces TLS in vivo. a, H&E staining of tumor sections from combination treatment groups. b, Multiplex immunofluorescence staining for CD3, CD19, CD23, and CD31 on tumor sections from the combination treatment group. c, Multiplex immunofluorescence staining for CD138, IgG and IgA on tumor sections from vehicle and combination treatment groups. d, Quantification of the CD138+ cell / mm^2^ and IgG/IgA ratio in tumor sections from vehicle, single-agent, and combination treatment groups. n = 3 per group. e, UMAP plots of B cells at indicated time points post-treatment. f, BCR repertoire diversity, clonal dynamics, and SHM following combination treatment. g, Immunoglobulin isotype distribution of B cells over the treatment time course. h, Peripheral BCR repertoire diversity and SHM following combination treatment. Statistical significance was determined by one-way ANOVA test. *P < 0.05; **P < 0.01; ***P < 0.001; ****P < 0.0001.

### Combination treatment elicits a B-cell-dependent in situ and memory antitumor effect

We next designed an experiment to dissect the B-cell-dependent in situ and memory antitumor effects of the combination regimen (Figure 7a). B cell depletion substantially abrogated the antitumor effect of the combination regimen, whereas depletion of CD4+, CD8+, or NK cells, or FcyR blockade, exerted minimal or no impact on therapeutic efficacy (Figure 7b and sFigure 6a). As humoral immunity primarily relies on antibody-dependent cellular cytotoxicity (ADCC), antibody-dependent cellular phagocytosis (ADCP), and complement-dependent cytotoxicity (CDC), we performed multiplex immunofluorescence staining for C3d, IgG, and CK8/18 to assess complement deposition and antibody binding on tumor cells. Given that neither FcyR receptor blockade nor NK depletion did not compromise treatment efficacy, ADCC and ADCP were not considered as the primary mechanisms underlying the observed antitumor effect. Diffuse C3d deposition was observed in tumors from the combination treatment group (Figure 7c). Furthermore, colocalization of CK8/18, C3d, and IgG was detected on tumor cell membranes (Figure 7c).

**Figure 7.**
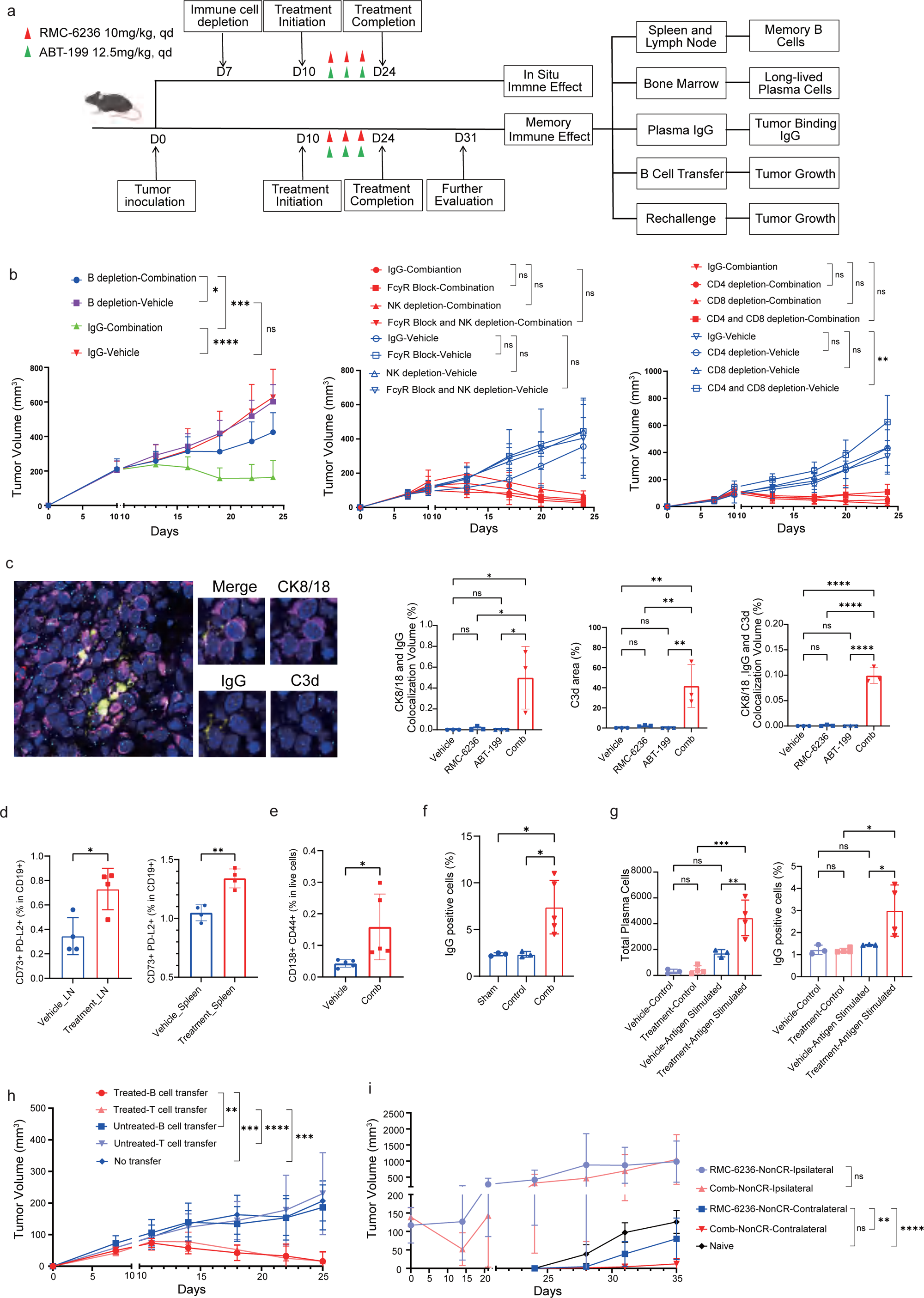
B-cell-dependent in situ and memory antitumor effects of the combination regimen. a, Experimental schematic of the in vivo depletion and adoptive transfer study design. b, Tumor growth curves of mice treated with the combination regimen following depletion of B cells, CD4+ T cells, CD8+ T cells, NK cells, or FcyR blockade. n = 5-8 mice per group. c, Multiplex immunofluorescence staining for C3d, IgG, and CK8/18 on tumor sections from vehicle, single-agent, and combination treatment groups. n = 3 per group. d, Quantification of memory B cells in the spleen and tumor-draining lymph nodes from vehicle and combination treatment groups by flow cytometry. n = 4 per group. e, Quantification of long-lived plasma cells in the bone marrow from vehicle and combination treatment groups by flow cytometry. n = 5 per group. f, Serum levels of tumor-specific antibodies measured by flow cytometry in vehicle and combination treatment groups. n = 3-5 per group. g, (Left) Quantification of plasma cells generated from CD19+ splenocytes stimulated with tumor antigens in vitro. (Right) IgG binding to tumor cells detected by flow cytometry after incubation with culture supernatants from vehicle or combination treatment groups. h, Tumor volume curves of naive recipient mice receiving adoptive transfer of B cells or T cells from vehicle- or combination-treated donor mice, followed by tumor inoculation and half-dose RMC-6236 administration. n = 7 mice per group. i, Rechallenge tumor growth in non-CR and treatment-naive mice. n = 8 mice per group. Statistical significance was determined by one-way ANOVA test. *P < 0.05; **P < 0.01; ***P < 0.001; ****P < 0.0001.

Memory B cells and long-lived plasma cells constitute the cellular basis of humoral immune memory. To characterize the memory response induced by the combination regimen, we performed a series of experiments. First, we observed an increase in memory B cells within the spleen and tumor-draining lymph nodes (Figure 7d, sFigure 6b and sFigure 6d), accompanied by long-lived plasma cells detected in the bone marrow (Figure 7e). Furthermore, elevated tumor-specific antibodies were detected in serum (Figure 7f), suggesting that memory B and long-lived plasma cells serve as the primary mediators of immune memory in this context.

To functionally validate memory B cell responses, we isolated CD19+ cells from spleens of treated mice and stimulated them with tumor antigens in vitro. Indeed, a greater number of plasma cells were generated from the combination group (Figure 7g and sFigure 6c). Consistently, incubation with the culture supernatants from the combination group resulted in increased IgG binding to tumor cells (Figure 7g). More importantly, we adoptively transferred B cells or T cells from vehicle- or combination-treated donor mice into naïve recipients. Following tumor inoculation, a half-dose of RMC-6236 was administered every other day to promote tumor antigen release while avoiding overt antitumor effects. Although both B cell and T cell recipients exhibited significant tumor shrinkage, B cell transfer induced a more rapid reduction in tumor volume (Figure 7h and sFigure 6e).

Finally, mice bearing small tumors (approximately 100 mm^3^) were treated with either RMC-6236 alone or the combination regimen for two weeks. Notably, 50% of mice in the monotherapy group and 80% in the combination group achieved complete response (CR), confirmed ten days following drug withdrawal. These mice were subsequently rechallenged with tumor cells on the contralateral flank to assess memory immunity for two weeks. No tumor growth was observed at the rechallenge site in any of the CR mice, regardless of the initial treatment. In contrast, among non-CR mice, the combination group exhibited significantly smaller tumor volumes in the contralateral flank, with 37.5% (3/8) remaining relapse-free compared with 0% (0/8) in the monotherapy group (Figure 7i and sFigure 6f).

In summary, the combination regimen established B-cell-mediated antitumor immunity, characterized by in situ CDC activity and durable memory responses through memory B cells and long-lived plasma cells. Adoptive transfer and rechallenge experiments confirmed that B cells are sufficient for rapid tumor control and long-term protection.

## Discussion

The biochemical and signaling heterogeneity among KRAS codon 12 mutations has been increasingly recognized over the past decade, with distinct alleles exhibiting differential effector engagement and pathway activation[4, 33]. However, whether these allele-specific features translate into distinct metabolic vulnerabilities and therapeutic opportunities remains incompletely understood. Our study extends this understanding by demonstrating that the G12D mutation exhibits a distinct mitochondrial vulnerability, characterized by weakened HK1 interaction, reduced spare respiratory capacity, and increased mitochondrial stress sensitivity. In the context of KRAS/STK11 co-mutation (KL), however, G12D is further associated with poor prognosis and a suppressed immune microenvironment, which we traced to reduced mitochondrial DNA leakage and subsequent silencing of type I interferon signaling, accompanied by impaired antigen presentation. These findings highlight the importance of integrating both mutation subtype and co-mutation status when dissecting the biology of KRAS-driven tumors, while also revealing that allele-specific mitochondrial vulnerabilities can exist independently of the co-mutational landscape.

We further found that this allele-specific mitochondrial vulnerability can be therapeutically exploited. Through a focused screen of clinically available agents that induce mitochondrial stress, we identified that combined inhibition of KRAS and BCL-2 synergistically triggers GSDME-mediated pyroptosis in G12D-mutant cells. Mechanistically, the combination of KRAS and BCL-2 inhibition led to a pronounced decline in mitochondrial membrane potential, accompanied by caspase-3 activation and GSDME cleavage—indicative of coordinated metabolic compromise and mitochondrial outer membrane permeabilization. This effect was specific to G12D-mutant cells, supporting the notion that allele-specific metabolic states can create context-dependent therapeutic windows.

Remarkably, this combination regimen initiates a cascade of antitumor immune remodeling. We observed the de novo formation of functional tertiary lymphoid structures (TLS)—ectopic lymphoid aggregates that support local humoral immune responses[34–36]. Within TLS, B cells underwent germinal center reactions, including affinity maturation and class switching, generating high-affinity IgG antibodies[37]. Notably, we detected C3d and IgG co-deposition on tumor cell membranes, while Fcγ receptor blockade did not compromise treatment efficacy—collectively indicating that CDC serves as the predominant effector mechanism of B-cell-mediated tumor killing in this context. Our findings thus establish a direct mechanistic link between immunogenic cell death and humoral immunity, expanding the current understanding of how immunogenic cell death modalities can engage adaptive immune responses.

Consistent with this model, immune cell depletion experiments demonstrated that B cells are indispensable for the therapeutic effect of the combination regimen: B-cell depletion largely abrogated antitumor efficacy, whereas depletion of CD4⁺ T cells, CD8⁺ T cells, or NK cells had minimal impact. Adoptive transfer of B cells from combination-treated donors conferred superior tumor control compared with T-cell transfer, indicating that B cells are not only necessary but also sufficient to mediate antitumor protection[38, 39]. Moreover, among mice that did not achieve complete response, combination-treated animals exhibited significantly superior memory protection upon rechallenge, with 37.5% (3/8) remaining tumor-free compared with 0% (0/8) in the monotherapy group, and significantly smaller tumor volumes overall. By contrast, all mice that achieved CR—regardless of treatment group—uniformly rejected rechallenge. These findings suggest that the combination regimen can establish durable immunological memory even in the absence of complete tumor regression, a scenario more representative of the clinical reality where most patients do not achieve CR, and highlight its potential to confer lasting antitumor immunity. These data position B-cell-mediated immunity as a central pillar of the long-term efficacy of this regimen.

Several limitations warrant consideration. First, while our data establish a phenotypical link between combination-induced pyroptosis and TLS formation. The precise molecular cascade linking pyroptosis-derived DAMPs to subsequent TLS neogenesis remains incompletely understood. Second, although we validated this strategy in multiple models, the safety and efficacy of combining KRAS and BCL-2 inhibitors in humans will require evaluation in clinical trials, which we view as a priority given the clinical availability of both agents. Despite these limitations, our findings provide a mechanistic framework and a strong translational rationale for targeting G12D-specific mitochondrial–immune crosstalk in KRAS G12D-mutant lung cancer.

## Materials and Methods

### Clinical Cohorts and Data Analysis

Clinical and gene mutation data were retrieved from the published pan-cancer cohort[12] and the lung cancer cohort treated with immunotherapy[13]. Transcriptomic data from TCGA (n = 86), GSE72094 (n = 78), and CCLE (n = 10) cohorts were analyzed. In the TCGA and GSE72094 cohorts, KL tumors were defined as KRAS/STK11 co-mutant or KRAS-mutant tumors with low STK11 expression. Low STK11 expression was defined as expression levels below the median level. Our single-center cohort data were obtained from the electronic medical record system.

Overall survival (OS) and progression-free survival (PFS) were analyzed using the Kaplan-Meier method, and survival differences between groups were compared using the log-rank test. ESTIMATE and IPS scores were calculated based on published algorithms[40, 41]. Gene set variation analysis (GSVA) was performed using the clusterProfiler package in R[42]. Gene set enrichment analysis (GSEA) was performed for pathway enrichment analysis based on Hallmark gene sets from the Molecular Signatures Database (MSigDB)[43].

### Cell Lines and Culture Conditions

Human lung cancer cell lines (A427, H2030, H23, H157, SKLU-1, H358, H441, H1792 and BEAS-2B), transgenic mouse-derived cell lines (KP-1, FC1242) (gifts from Prof. Fei Li from School of Basic Medical Sciences, Fudan University and Prof. Shengxiang Ren from Shanghai Pulmonary Hospital, Tongji University School of Medicine) and mouse-derived cell lines (CT26) were cultured in RPMI-1640 or DMEM supplemented with 10% fetal bovine serum (FBS), 100 U/mL penicillin, and 100 μg/mL streptomycin at 37°C in 5% CO_2_. All cell lines were routinely tested for mycoplasma contamination.

### Transgenic Mouse Models

Kras^LSL-G12D/+ and Kras^LSL-G12C/+ mice were crossed with Stk11^f/f mice to generate Kras^G12D/+;Stk11^f/f and Kras^G12C/+;Stk11^f/f mice. All mice were maintained on a C57BL/6 background. To induce Cre-mediated recombination and activate oncogenic Kras expression while knocking out Stk11, eight to twelve-week-old mice were anesthetized, and intranasal inhalation of Cre-expressing adenovirus (CMV-Cre) was performed as described[44]. Briefly, 2*10⁶ PFU of CMV-Cre was mixed with 60.2 μL of MEM and 0.3 μL of CaCl₂ to a final volume of 62.5 μL per mouse. Ten weeks later, mice were sacrificed for further experiments.

### Organoid Culture from Transgenic Mice

Lung tumors from Kras^G12D/+;Stk11^f/f (KDL) and Kras^G12C/+;Stk11^f/f (KCL) mice were dissected, minced, and digested in DMEM containing collagenase IV (1 mg/mL) and DNase I (0.1 mg/mL) for 30 minutes at 37°C.Dissociated cells were embedded in growth factor-reduced Matrigel (Corning) and cultured in organoid growth medium (JiaShiTeng, China). Organoids were passaged every 7-10 days and used for experiments between passages.

### Subcutaneous Xenograft Models

KP-1 and FC1242 cells (1*10^6^) were suspended in 100 μL of PBS and injected subcutaneously into the right flank of 5-6 week old female immunocompromised mice. When tumors reached approximately 200 ml, mice were randomized into treatment groups. RMC-6236 (10 mg/kg) and ABT-199 (12.5 mg/kg) were administered orally once daily. Tumor volumes were measured twice a week using a caliper.

For re-challenge experiments, the same amount of cells were injected subcutaneously into the left flank of the treated mice ten days after two weeks of treatment.

For adoptive transfer experiments, 2 × 10⁶ isolated B or T cells from the spleens of donor mice were injected via tail vein into recipient mice. The following day, recipient mice were inoculated subcutaneously with 1 × 10⁶ KP-1 tumor cells. Ten days after tumor inoculation, a half dose of RMC-6236 (5 mg/kg) was administered every other day, and tumor volumes were measured twice a week thereafter.

### Co-immunoprecipitation (Co-IP)

For Co-IP, cell lysates were incubated with anti-FLAG or anti-EGFP Nanoab-Magnetic Beads (LABLEAD) at 4°C for 60 min. Beads were washed three times, and bound proteins were eluted by boiling in SDS sample buffer for immunoblot analysis.

### Fluorescence Microscopy

For JC-1 or staining, cells were incubated with JC-1 dye for 20 minutes at 37°C, washed twice with PBS, and immediately imaged using a confocal microscope (Zeiss LSM 880). Images were acquired with a 10* objective lens.

For cytosolic mtDNA detection, cells were first incubated with MitoTracker (200 nM) for 40 minutes at 37°C to label mitochondria, followed by staining with the mtDNA-specific probe PicoGreen for 20 minutes. Images were acquired using a confocal microscope (Zeiss LSM 880) with a 63* oil immersion objective.

For mitochondrial morphology analysis, cells were stained with MitoTracker (1:2,500 dilution in Hank’s balanced salt solution) for 10 minutes at 37°C. Cells were washed twice with PBS and imaged in Hank’s solution using a super-resolution live-cell imaging system (OMX SR).

### Flow Cytometry

For MMP measurement, cells were stained with JC-1 dye for 20 minutes at 37°C, washed, and analyzed. MMP was quantified as the ratio with high red (JC-1 aggregates, 580 nm) fluorescence. For apoptosis detection, cells were stained with Annexin V and 7-AAD for 15 minutes according to the manufacturer’s protocol (MULTI SCIENCES, AP104).

For immune cell profiling, tumor tissues were dissociated into single-cell suspensions. Cells were stained with ZOMBIE dye for viability assessment, followed by Fc receptor blockade with anti-CD16/32 antibody (clone S17011E). Surface and intracellular staining was then performed using antibodies against CD45 (clone I3/2.3), CD3 (clone 17A2), CD4 (clone RM4-5), CD8 (clone 53-6.7), CD44 (clone IM7), CD19 (clone 6D5), CD49b (clone DX5), GzmB (clone QA16A02), IFN-γ (clone XMG1.2), CD11c (clone N418), CD86 (clone GL-1), CD206 (clone M5/114.15.2), CD103 (clone 2E7), F4/80 (clone BM8), CD11b (clone M1/70) and Gr-1 (clone RB6-8C5). All antibodies were used according to the manufacturer’s protocols (Biolegend).

Data were acquired on a flow cytometer (SONY ID7000) and analyzed using FlowJo software.

### Immune Cell Depletion

Mice were injected intraperitoneally with monoclonal antibodies (200 μg/mouse) targeting CD4 (clone GK1.5, BioXCell BE0003-1), CD8 (clone 53-6.7, BioXCell BE0003-1), or NK cells (clone PK136, BioXCell BE0036) twice weekly. For B cell depletion, anti-B cell antibody (clone MB20-11, BioXCell BP0356) was administered once via tail vein injection at 10 μg per mouse. For Fcγ receptor blockade, mice received anti-FcγR antibody (clone 9E9, BioXCell BE0378) at 200 μg/mouse twice weekly. Isotype-matched control antibodies were used as negative controls. Depletion was initiated three days prior to treatment start. Depletion efficiency was confirmed by flow cytometry analysis of peripheral blood lymphocytes.

### Tumor Binding Immunoglobin Detection Assay

For tumor-binding Ig detection, harvested tumor cells (5 × 10⁵ per sample) were blocked with 1% BSA/PBS for 10 minutes at 4°C. Cells were then incubated with mouse plasma (1:5 dilution in PBS) for 60 minutes at 4°C. After washing, bound antibodies were detected with anti-IgG-AF647 and anti-IgM-AF488 (1:2000 each) for 30 minutes at 4°C, followed by 7-AAD staining for viability assessment prior to flow cytometry.

### Splenic B and T cell Isolation

Splenocytes from vehicle- or combination-treated mice were pooled (2 spleens per group) and processed into single-cell suspensions using a lymphocyte separation kit (Solarbio, P8860). After density gradient centrifugation (800g, 30 min), lymphocytes were collected from the interface. CD19+ B cells were positively selected using CD19 microbeads (Miltenyi Biotec, 130-121-301) following the manufacturer’s protocol. Briefly, cells were incubated with CD19 microbeads (40 μL per group) for 20 minutes at 4°C, then separated on magnetic columns. The CD19-cells was collected, and CD19+ cells were eluted from the columns. The CD19-fraction was then subjected to CD3+ T cell isolation using negative selection microbeads following the manufacturer’s protocol (Selleck, B90021).

### Tumor Antigen Preparation

Tumor cells were trypsinized, washed twice with PBS (400g, 5 min), and resuspended in PBS at 1*10⁷ cells/mL. Cell suspensions were transferred to cryotubes and subjected to three cycles of freeze-thaw lysis: freezing in liquid nitrogen (1-2 min) followed by thawing in a 37°C water bath. After the final thaw, lysates were centrifuged at 10000g for 15 min at 4°C. The supernatant was collected, filtered through a 0.22 μm filter, and stored at −80°C until use.

### Memory B Cell in Vitro Stimulation

Isolated splenic CD19+ B cells (2*10⁶ cells per well) were cultured in 1 mL of conditioned medium containing 33 μL tumor antigen, IL-2 (100 IU/mL), IL-21 (50 ng/mL), CpG (0.25 μg/mL), 10% FBS, and antibiotics for 48 hours. Cells were then harvested and analyzed for plasma cell differentiation by flow cytometry.

## References

1. Kris, M.G., et al., Using multiplexed assays of oncogenic drivers in lung cancers to select targeted drugs. Jama, 2014. 311(19): p. 1998–2006.

2. Ricciuti, B., et al., Dissecting the clinicopathologic, genomic, and immunophenotypic correlates of KRAS(G12D)-mutated non-small-cell lung cancer. Ann Oncol, 2022. 33(10): p. 1029–1040.

3. Skoulidis, F., et al., Co-occurring genomic alterations define major subsets of KRAS-mutant lung adenocarcinoma with distinct biology, immune profiles, and therapeutic vulnerabilities. Cancer Discov, 2015. 5(8): p. 860–77.

4. Hunter, J.C., et al., Biochemical and Structural Analysis of Common Cancer-Associated KRAS Mutations. Mol Cancer Res, 2015. 13(9): p. 1325–35.

5. Ihle, N.T., et al., Effect of KRAS oncogene substitutions on protein behavior: implications for signaling and clinical outcome. J Natl Cancer Inst, 2012. 104(3): p. 228–39.

6. Varshavi, D., et al., Metabolic characterization of colorectal cancer cells harbouring different KRAS mutations in codon 12, 13, 61 and 146 using human SW48 isogenic cell lines. Metabolomics, 2020. 16(4): p. 51.

7. Nuevo-Tapioles, C., et al., KRAS4A promotes oligomerization of hexokinase 1 on mitochondria. Cell Rep, 2026. 45(6): p. 117392.

8. Amendola, C.R., et al., KRAS4A directly regulates hexokinase 1. Nature, 2019. 576(7787): p. 482–486.

9. Cook, J.H., et al., The origins and genetic interactions of KRAS mutations are allele- and tissue-specific. Nat Commun, 2021. 12(1): p. 1808.

10. Koyama, S., et al., STK11/LKB1 Deficiency Promotes Neutrophil Recruitment and Proinflammatory Cytokine Production to Suppress T-cell Activity in the Lung Tumor Microenvironment. Cancer Res, 2016. 76(5): p. 999–1008.

11. Skoulidis, F., et al., STK11/LKB1 Mutations and PD-1 Inhibitor Resistance in KRAS-Mutant Lung Adenocarcinoma. Cancer Discov, 2018. 8(7): p. 822–835.

12. Nguyen, B., et al., Genomic characterization of metastatic patterns from prospective clinical sequencing of 25,000 patients. Cell, 2022. 185(3): p. 563–575.e11.

13. Samstein, R.M., et al., Tumor mutational load predicts survival after immunotherapy across multiple cancer types. Nature Genetics, 2019. 51(2): p. 202–206.

14. Schabath, M.B., et al., Differential association of STK11 and TP53 with KRAS mutation-associated gene expression, proliferation and immune surveillance in lung adenocarcinoma. Oncogene, 2016. 35(24): p. 3209–16.

15. Decout, A., et al., The cGAS-STING pathway as a therapeutic target in inflammatory diseases. Nat Rev Immunol, 2021. 21(9): p. 548–569.

16. Green, D.R. and F. Llambi, Cell Death Signaling. Cold Spring Harb Perspect Biol, 2015. 7(12).

17. Liu, Y., et al., Understanding the complexity of p53 in a new era of tumor suppression. Cancer Cell, 2024. 42(6): p. 946–967.

18. Taniguchi, K. and M. Karin, NF-κB, inflammation, immunity and cancer: coming of age. Nat Rev Immunol, 2018. 18(5): p. 309–324.

19. Zhang, Y., et al., LKB1 deficiency-induced metabolic reprogramming in tumorigenesis and non-neoplastic diseases. Mol Metab, 2021. 44: p. 101131.

20. Hunter, J.C., et al., Biochemical and Structural Analysis of Common Cancer-Associated KRAS Mutations. Molecular Cancer Research, 2015. 13(9): p. 1325–1335.

21. Nickens, K.P., et al., A bioenergetic profile of non-transformed fibroblasts uncovers a link between death-resistance and enhanced spare respiratory capacity. Mitochondrion, 2013. 13(6): p. 662–7.

22. Azoulay-Zohar, H., et al., In self-defence: hexokinase promotes voltage-dependent anion channel closure and prevents mitochondria-mediated apoptotic cell death. Biochem J, 2004. 377(Pt 2): p. 347–55.

23. Houghton, J.A., et al., Inhibition of apoptosis after thymineless stress is conferred by oncogenic K-Ras in colon carcinoma cells. Clin Cancer Res, 1998. 4(11): p. 2841–8.

24. Bryant, K.L., et al., Combination of ERK and autophagy inhibition as a treatment approach for pancreatic cancer. Nat Med, 2019. 25(4): p. 628–640.

25. Vogler, M., et al., The BCL2 family: from apoptosis mechanisms to new advances in targeted therapy. Signal Transduct Target Ther, 2025. 10(1): p. 91.

26. Behranvand, N., et al., Chemotherapy: a double-edged sword in cancer treatment. Cancer Immunol Immunother, 2022. 71(3): p. 507–526.

27. Osman, A.M.A. and A.A. Arabi, Targeting mitochondrial complexes for cancer therapy. Biochem Pharmacol, 2026. 247: p. 117695.

28. Wang, Y., et al., Chemotherapy drugs induce pyroptosis through caspase-3 cleavage of a gasdermin. Nature, 2017. 547(7661): p. 99–103.

29. Lu, H., et al., Molecular Targeted Therapies Elicit Concurrent Apoptotic and GSDME-Dependent Pyroptotic Tumor Cell Death. Clin Cancer Res, 2018. 24(23): p. 6066–6077.

30. Erkes, D.A., et al., Mutant BRAF and MEK Inhibitors Regulate the Tumor Immune Microenvironment via Pyroptosis. Cancer Discovery, 2020. 10(2): p. 254–269.

31. Ma, J., et al., A blueprint for tumor-infiltrating B cells across human cancers. Science, 2024. 384(6695).

32. Zhang, Y., et al., CCL19-producing fibroblasts promote tertiary lymphoid structure formation enhancing anti-tumor IgG response in colorectal cancer liver metastasis. Cancer Cell, 2024. 42(8): p. 1370–1385.e9.

33. Simanshu, D.K. and F. McCormick, Revisiting RAS family GTPase signaling: effector selectivity and oncogenic bypass. Biochem J, 2026. 483(5): p. 761–775.

34. Sautès-Fridman, C., et al., Tertiary Lymphoid Structures and B cells: Clinical impact and therapeutic modulation in cancer. Semin Immunol, 2020. 48: p. 101406.

35. Schumacher, T.N. and D.S. Thommen, Tertiary lymphoid structures in cancer. Science, 2022. 375(6576).

36. Dieu-Nosjean, M.C., et al., Tertiary lymphoid structures in cancer and beyond. Trends Immunol, 2014. 35(11): p. 571–80.

37. Rodriguez, A.B., et al., Immune mechanisms orchestrate tertiary lymphoid structures in tumors via cancer-associated fibroblasts. Cell Rep, 2021. 36(3): p. 109422.

38. Glaros, V., N. Francis, and T. Kreslavsky, The multilayered identity of B cell memory. Cell Mol Immunol, 2026. 23(2): p. 150–167.

39. Kuan, V.L.S. and J. Chaudhuri, Mechanisms Promoting Stability of B Cells. Immunol Rev, 2025. 336(1): p. e70064.

40. Yoshihara, K., et al., Inferring tumour purity and stromal and immune cell admixture from expression data. Nat Commun, 2013. 4: p. 2612.

41. Charoentong, P., et al., Pan-cancer Immunogenomic Analyses Reveal Genotype-Immunophenotype Relationships and Predictors of Response to Checkpoint Blockade. Cell Rep, 2017. 18(1): p. 248–262.

42. Hänzelmann, S., R. Castelo, and J. Guinney, GSVA: gene set variation analysis for microarray and RNA-seq data. BMC Bioinformatics, 2013. 14: p. 7.

43. Subramanian, A., et al., Gene set enrichment analysis: a knowledge-based approach for interpreting genome-wide expression profiles. Proc Natl Acad Sci U S A, 2005. 102(43): p. 15545–50.

44. DuPage, M., A.L. Dooley, and T. Jacks, Conditional mouse lung cancer models using adenoviral or lentiviral delivery of Cre recombinase. Nat Protoc, 2009. 4(7): p. 1064–72.

