## Supplementary figures and images for "Co-targeting KRAS and BCL-2 induces pyroptosis and antitumor tertiary lymphoid structures"

### Supplemental Figure 1

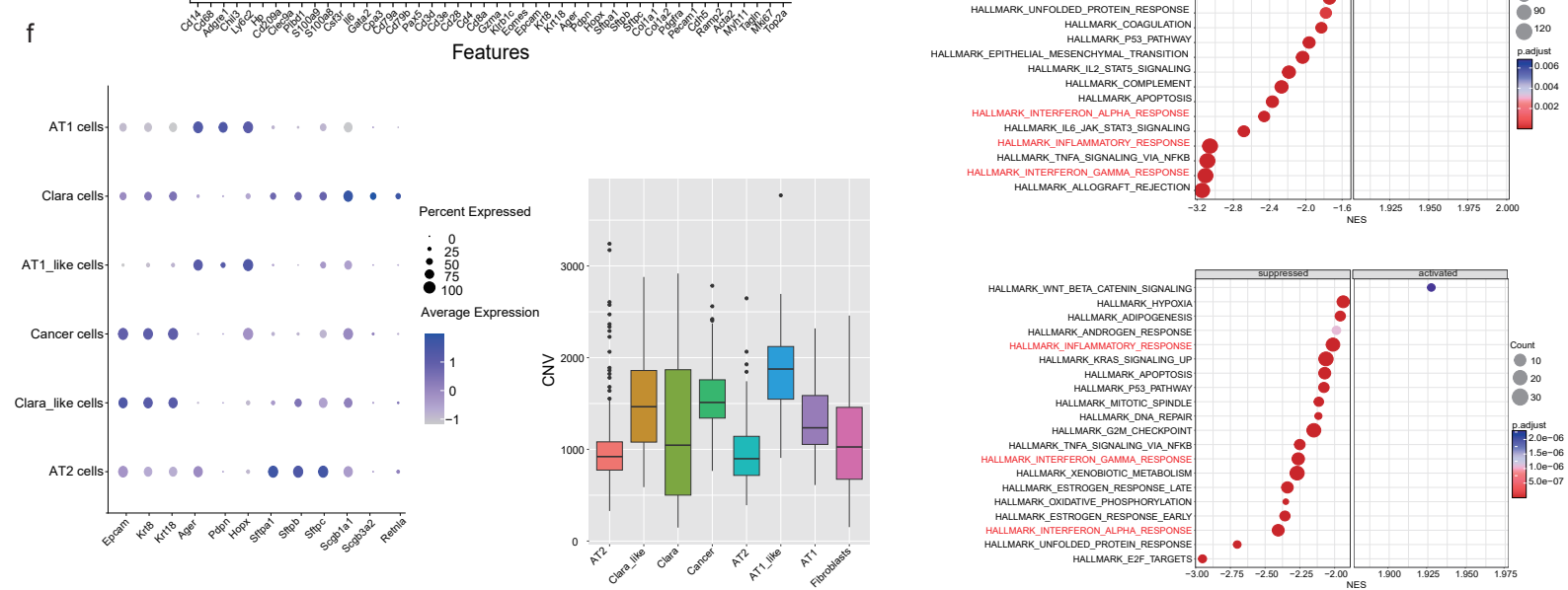

### Supplemental Figure 2

a

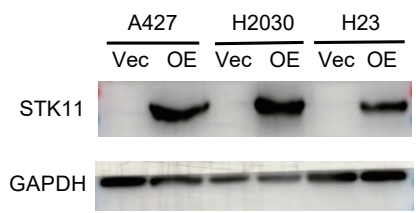

b

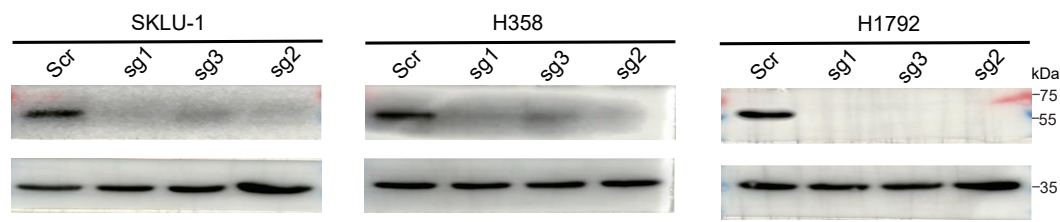

c

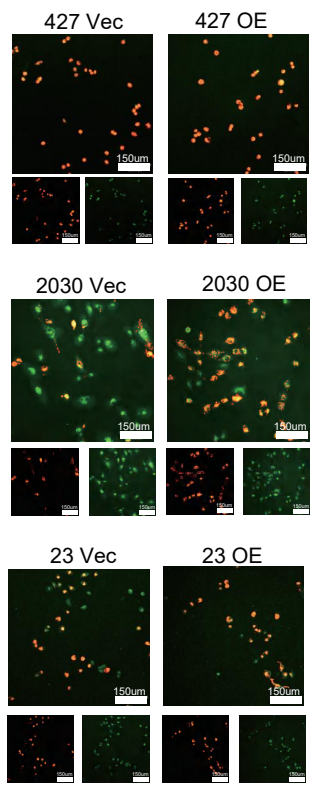

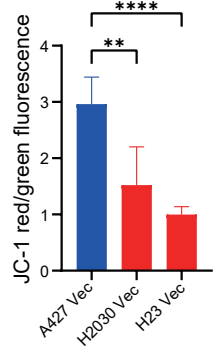

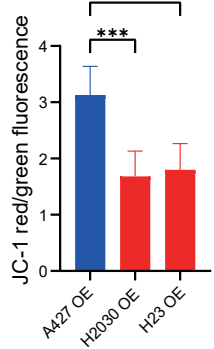

d

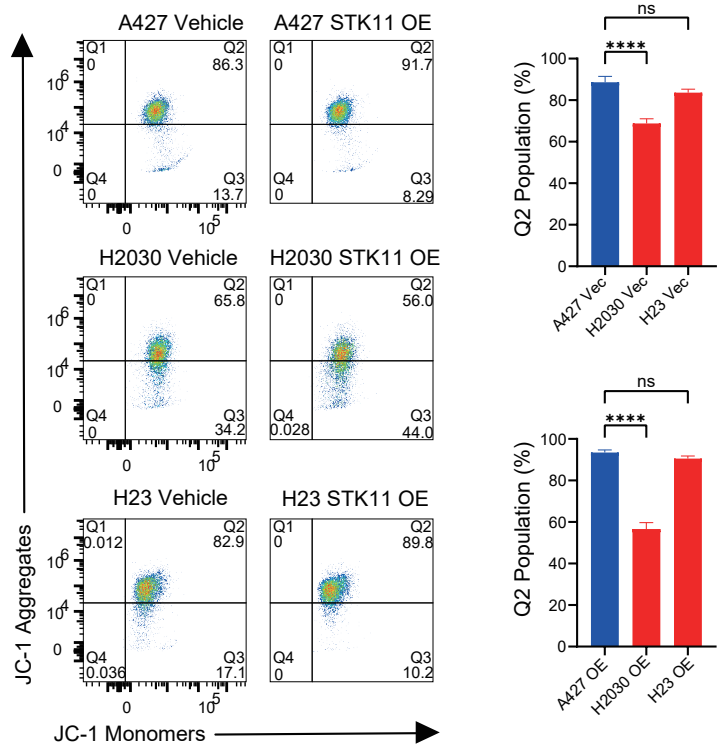

e

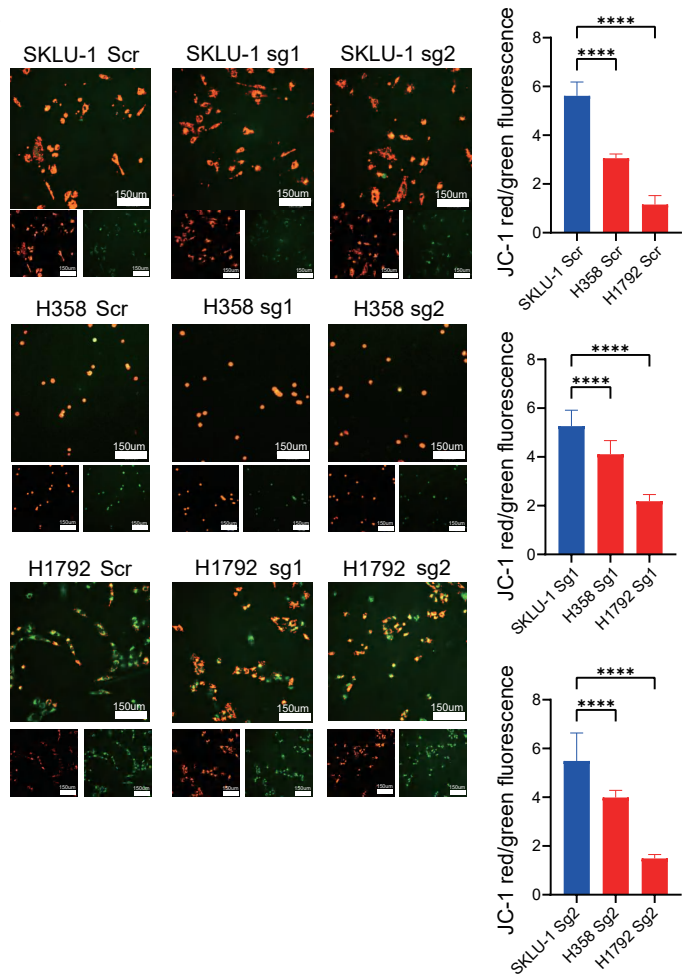

f

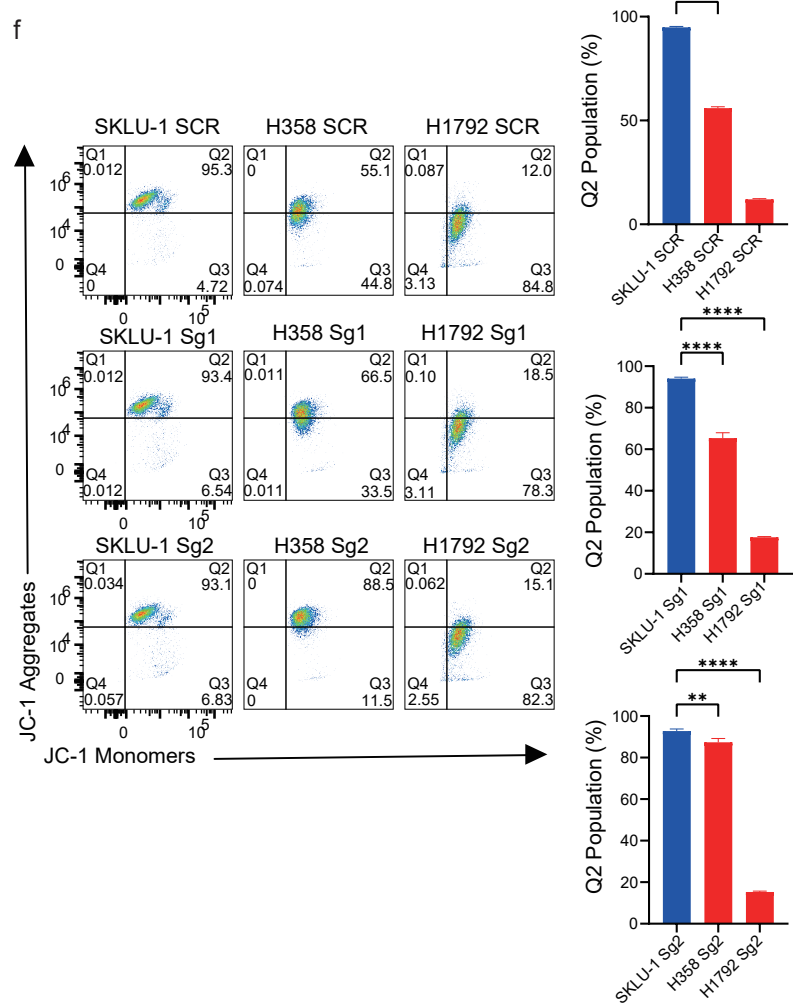

### Supplemental Figure 3

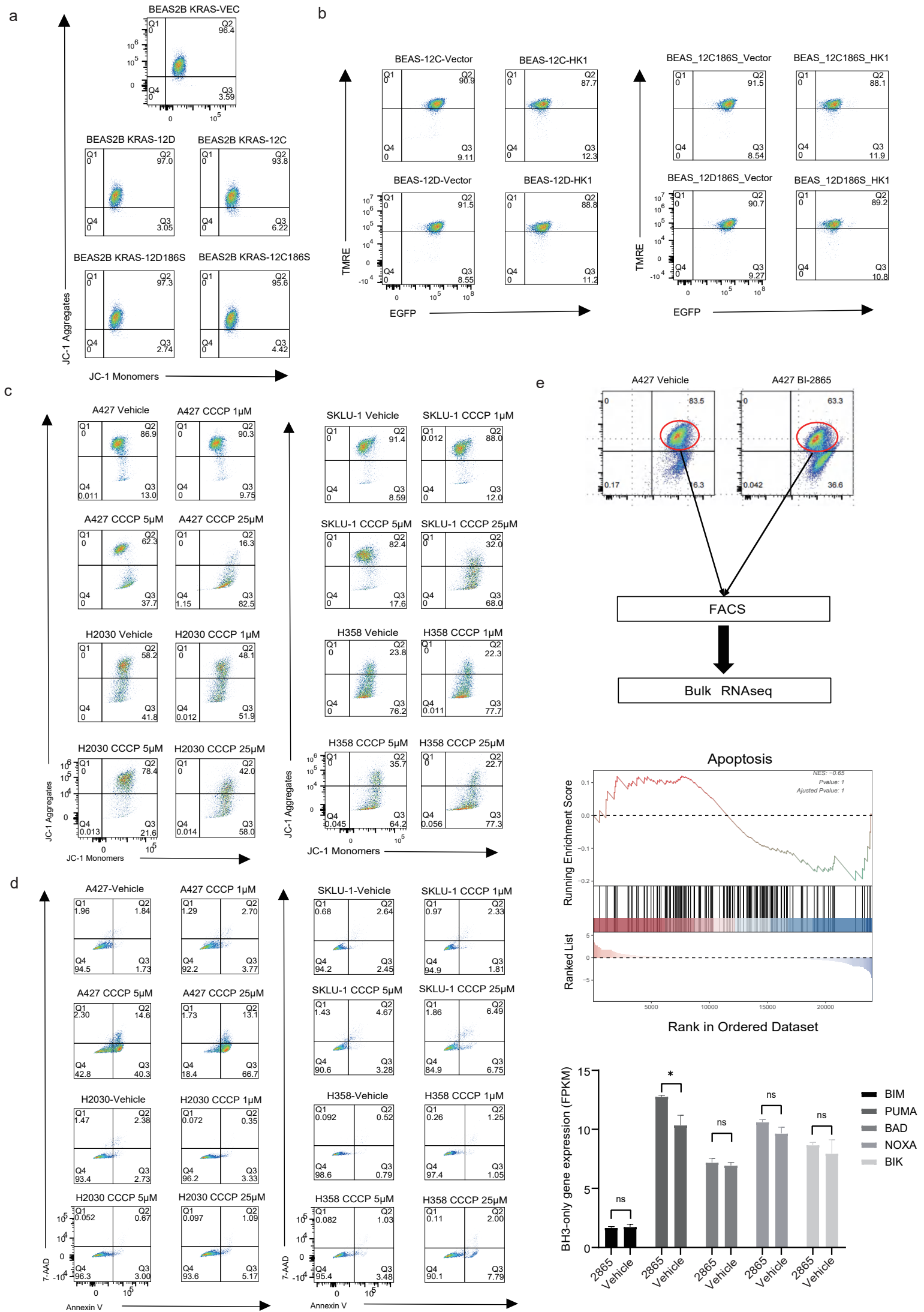

### Supplemental Figure 4

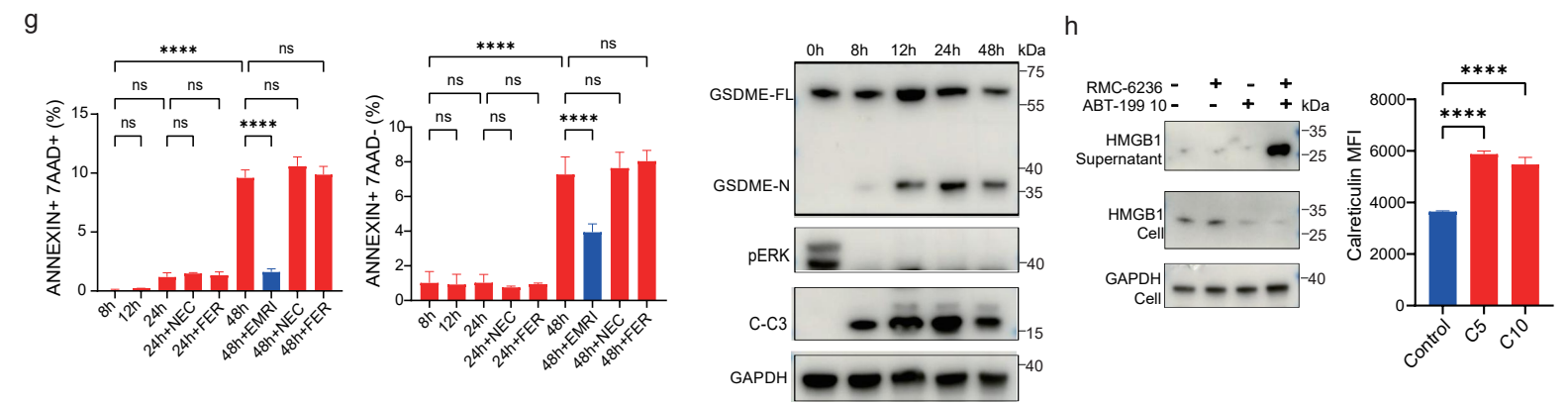

### Supplemental Figure 5

a

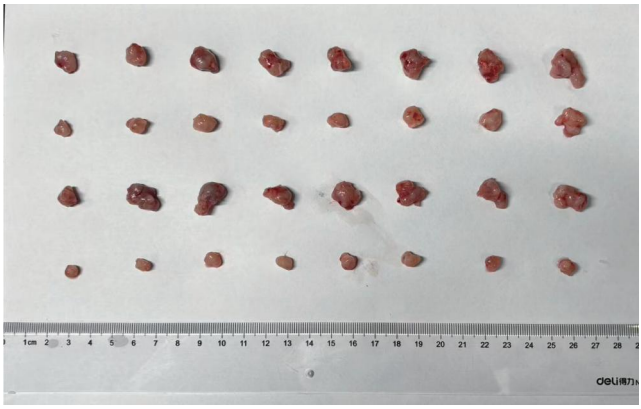

b

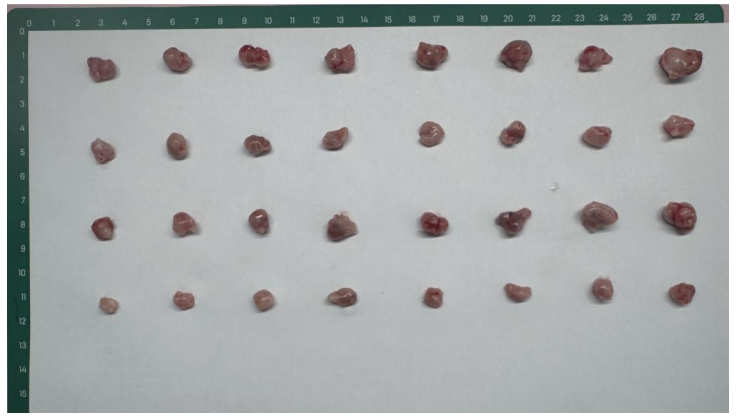

c

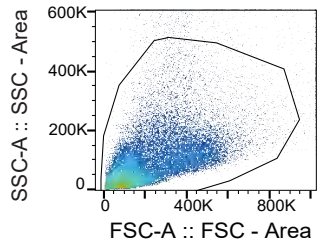

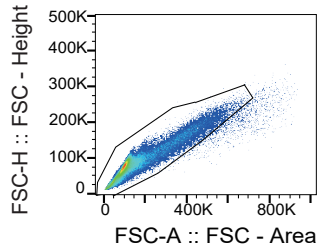

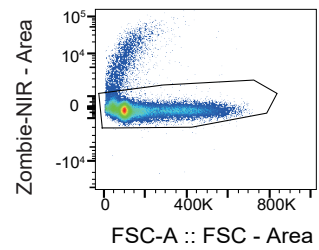

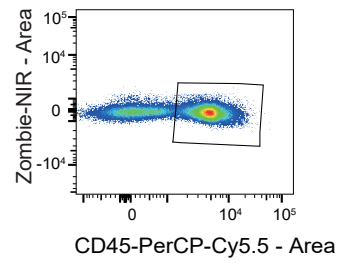

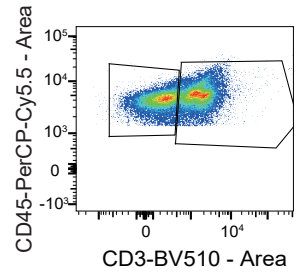

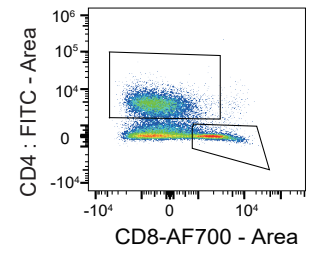

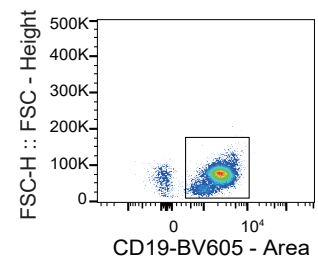

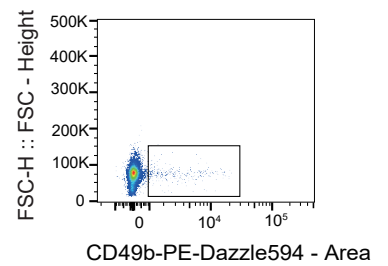

d

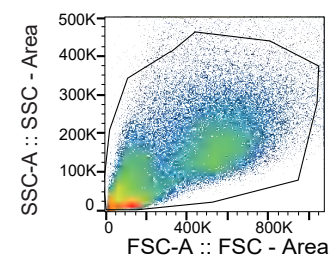

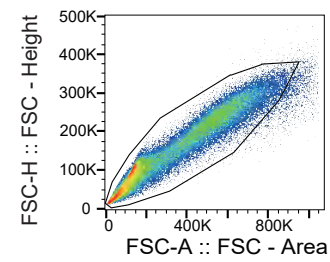

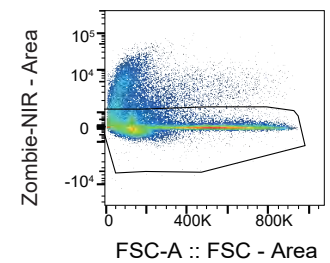

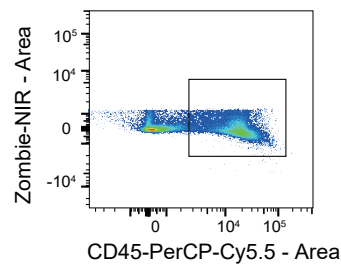

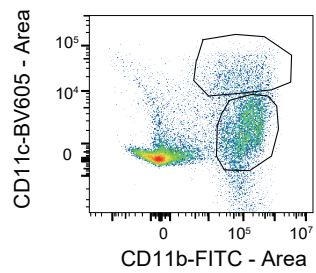

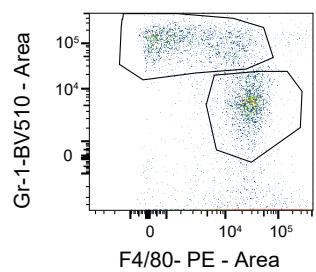

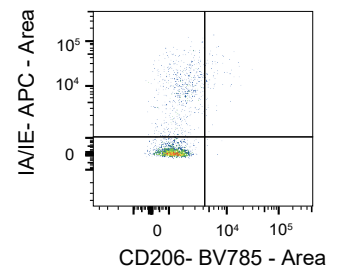

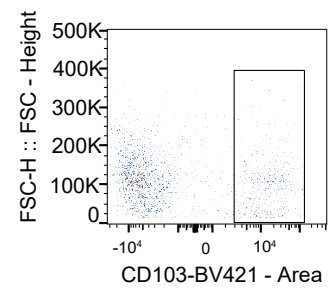

### Supplemental Figure 6

a

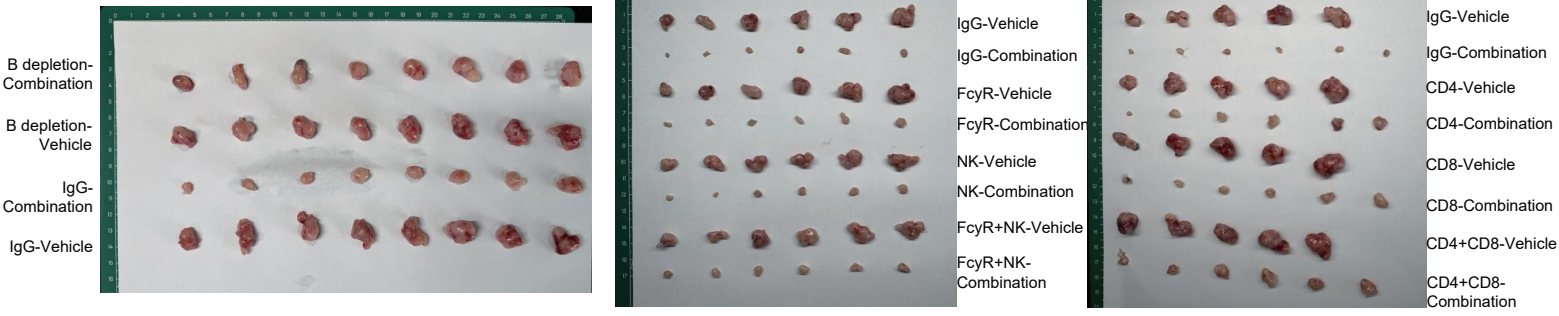

b

c

d

e

f
