## Supplementary material for "Co-targeting KRAS and BCL-2 induces pyroptosis and antitumor tertiary lymphoid structures": Figure Legend Supplement

**sFigure 1**

- a, b, Kaplan – Meier survival curves for overall survival (OS) in a published pan-cancer cohort (up) and an immunotherapy cohort (down), stratified by KRAS mutation and co-mutation subtypes;
- c, IPS and ESTIMATE analysis from TCGA dataset comparing G12D versus non-G12D tumors within STK11-low/co-mutant KL settings.
- d, Dotplots of single-cell RNA-seq data in KDL and KCL tumors.
- e, GSEA analysis of downregulated pathways in KRAS G12D versus non-G12D KL tumors from the TCGA (n = 86), GSE72094 (n = 78), and CCLE (n = 10) datasets.
- f, Dotplots and infer-CNV analysis of epithelium cells in KDL and KCL tumors.

**sFigure 2**

a, b, Western blot of STK11 overexpression and knock-out in KL and KP cells, respectively.

c, Representative confocal microscopy images of JC-1 staining in KL cells with STK11 overexpression. Red fluorescence indicates JC-1 aggregates; green fluorescence indicates JC-1 monomers. n = 3 biological replicates, each with 3 technical replicates.

d, Quantification of MMP by flow cytometry in KL cells with STK11 overexpression. n = 3 biological replicates, each with 3 technical replicates.

e, Representative confocal microscopy images of JC-1 staining in KP cells with STK11 knock-out. Red fluorescence indicates JC-1 aggregates; green fluorescence indicates JC-1 monomers. n = 3 biological replicates, each with 3 technical replicates.

f, Quantification of MMP by flow cytometry in KP cells with STK11 knock-out. n = 3 biological replicates, each with 3 technical replicates.

Statistical significance was determined by one-way ANOVA test. \*P < 0.05; \*\*P < 0.01; \*\*\*P < 0.001; \*\*\*\*P < 0.0001.

**sFigure 3.**

a, MMP in BEAS-2B cells expressing FLAG-tagged KRAS G12C, G12D, G12C186S and G12D186S mutants.

b, MMP analysis in BEAS-2B cells expressing FLAG-tagged KRAS mutants with or without HK1-EGFP overexpression.

c, d, MMP and apoptosis assays in G12D and G12C cells following CCCP treatment.

e, Bulk RNA sequencing of MMP-high versus MMP-low A427 cells following BI-2865 treatment. n = 3 biological replicates.

Western blots and co-IP were representative of three independent biological replicates. Statistical significance was determined by one-way ANOVA test. \*P < 0.05; \*\*P < 0.01; \*\*\*P < 0.001; \*\*\*\*P < 0.0001.

**sFigure 4.**

a, Synergistic effects of BI-2865 combined with BH3 mimetics, chemotherapeutic agents, or mitochondrial complex inhibitors in A427 cells.

b, Synergistic effect of combined BI-2865 and ABT-199 in transgenic mouse-derived organoids and human cell lines.

c, d, Flow cytometry analysis of apoptosis in KL and KP cell lines following combination treatment. n = 3 biological replicates, each with 3 technical replicates.

e, f, Flow cytometry analysis of MMP and apoptosis in mouse KP-1 and FC1242 cell lines.

g, Flow cytometry of apoptosis and immunoblot of cell death markers in A427 cell lines following combination treatment. n = 3 biological replicates, each with 3 technical replicates.

h, Cell supernatant HMGB1 (left) and Surface calreticulin (right) in A427 cell lines following treatment. n = 3 biological replicates, each with 3 technical replicates.

Western blots were representative of three independent biological replicates. Statistical significance was determined by one-way ANOVA test. \*P < 0.05; \*\*P < 0.01; \*\*\*P < 0.001; \*\*\*\*P < 0.0001.

**sFigure 5.**

a, b, Antitumor efficacy of combination treatment in KP-1 and FC1242 subcutaneous xenograft models. n = 8 mice per group.

c, Gating strategy for Lymphoid immune cell composition in KP1 tumors.

d, Gating strategy for Myeloid immune cell composition in KP1 tumors. n = 6 per group.

**sFigure 6**

a, Tumors treated with the combination regimen following depletion of B cells, CD4<sup>+</sup> T cells, CD8<sup>+</sup> T cells, NK cells, or FcγR blockade. n = 5-8 mice per group.

b, Gating strategy for memory B cells.

c, Gating strategy for plasma B cells.

d, B and T cell activation and effector/memory phenotypes in secondary lymphoid organs in vehicle and combination treatment groups. n = 4 per group.

e, Tumors of naive recipient mice receiving adoptive transfer of B cells or T cells from vehicle- or combination-treated donor mice, followed by tumor inoculation and half-dose RMC-6236 administration. n = 7 mice per group.

f, Rechallenge tumors in non-CR and treatment-naive mice. n = 8 mice per group. Statistical significance was determined by one-way ANOVA test. \*P < 0.05; \*\*P < 0.01; \*\*\*P < 0.001; \*\*\*\*P < 0.0001.

.
